# Targeting transcriptional coregulator OCA-B/Pou2af1 blocks activated autoreactive T cells in the pancreas and type-1 diabetes

**DOI:** 10.1101/2020.02.06.937839

**Authors:** Heejoo Kim, Jelena Perovanovic, Arvind Shakya, Zuolian Shen, Cody N. German, Andrea Ibarra, Jillian L. Jafek, Nai-Pin Lin, Brian D. Evavold, Danny H.-C. Chou, Peter E. Jensen, Xiao He, Dean Tantin

## Abstract

The transcriptional coregulator OCA-B promotes expression of T cell target genes in cases of repeated antigen exposure, a necessary feature of autoimmunity. We hypothesized that T cell-specific OCA-B deletion and pharmacologic OCA-B inhibition would protect mice from autoimmune diabetes. We developed an *Ocab* conditional allele and backcrossed it onto a diabetes-prone NOD/ShiLtJ strain background. T cell-specific OCA-B loss protected mice from spontaneous disease. Protection was associated with large reductions in islet CD8^+^ T cell receptor specificities associated with diabetes pathogenesis. CD4^+^ clones associated with diabetes were present, but associated with anergic phenotypes. The protective effect of OCA-B loss was recapitulated using autoantigen-specific NY8.3 mice, but diminished in monoclonal models specific to artificial or neoantigens. Rationally-designed membrane-penetrating OCA-B peptide inhibitors normalized glucose levels, and reduced T cell infiltration and proinflammatory cytokine expression in newly-diabetic NOD mice. Together, the results indicate that OCA-B is a potent autoimmune regulator and a promising target for pharmacologic inhibition.

**~40-word summary statement for the online JEM table of contents and alerts:** Kim and colleagues show that OCA-B in T cells is essential for the generation of type-1 diabetes. OCA-B loss leaves the pancreatic lymph nodes largely undisturbed, but associates autoreactive CD4^+^ T cells in the pancreas with anergy while deleting potentially autoreactive CD8^+^ T cells.

**Summary:** Kim et al. show that loss or inhibition of OCA-B in T cells protects mice from type-1 diabetes.

## Introduction

Type-1 diabetes (T1D) is an autoimmune disease in which the host immune system is directed towards antigens associated with pancreatic beta cells (Cooke and Plotnick, 2008; Lernmark and Larsson, 2013). Pathologically, T1D is characterized by insulitis, beta cell destruction and inability to produce insulin. The main treatment for T1D, life-long insulin therapy, treats symptoms but not cause. The development of new T1D treatments is limited by an incomplete understanding of disease mechanisms (Staeva et al., 2013). Beta cell regeneration is a promising line of therapy, but still requires methods to specifically block T1D autoimmunity. An ideal therapy would block autoimmunity early in the disease course to spare remaining beta cell function while preserving normal immune function.

OCA-B, also known as Bob.1/OBF-1 (gene symbol *Pou2af1*), is a transcriptional coregulatory protein named for its strong expression in the B cell lineage, where it is dispensable until after B cell activation. Following B cell activation, it is essential for the generation of germinal centers (Shi et al., 2015). In CD4^+^ T cells, OCA-B docks with the transcription factor Oct1 to regulate a set of ~150 target genes, including *Il2, Ifng* and *Csf2* (*Gmcsf*) (Shakya et al., 2015). Upon T cell activation, many of these targets are activated by pathways that converge on transcription factors such as NF-AT, AP-1 and NF-kB. Factors like NF-AT can be thought of as the primary on/off switches for these genes, and drugs that block their activity effectively block target gene expression. Such drugs have utility in many contexts, but also have drawbacks including global dampening of immune function and side effects due to expression in other tissues. In contrast, Oct1 and OCA-B insulate silent but previously activated target genes against stable repression. Loss of either protein does not affect CD4^+^ T cell responses to stimulation in vitro or primary infection in mice (Kim et al., 2019; Shakya et al., 2015), but causes target gene expression defects upon secondary stimulation in vitro (Shakya et al., 2015; Shakya et al., 2011). Loss of either protein also results in defective CD4^+^ memory T cell formation and recall responses in vivo (Shakya et al., 2015). In addition, OCA-B expression is largely confined to lymphocytes. OCA-B is undetectable in thymocytes and naïve CD4^+^ T cells, but becomes stably expressed after antigen stimulation (Shakya et al., 2015). Once expressed, OCA-B recruits the histone lysine demethylase Jmjd1a/Jhdm2a/Kdm3a to Oct1 at target loci, where it locally removes inhibitory H3K9me2 chromatin marks that would otherwise promote repressive chromatin environments (Shakya et al., 2015). OCA-B also potentiates polarization of activated helper T cells towards the Th17 phenotype, among the most pro-inflammatory helper T cell subsets (Yosef et al., 2013).

Persistent antigen exposure is a common property of autoimmune responses. Among profiled CD4^+^ T cell subsets, OCA-B levels are elevated in both central memory cells and pancreas-infiltrating, islet-reactive CD4^+^ T cells (Heng et al., 2008). Both Oct1 and OCA-B are highly conserved from mice to humans, and genome-wide association studies identify Oct1/OCA-B binding site polymorphisms that confer predisposition to a variety of human autoimmune diseases including T1D, multiple sclerosis, lupus, inflammatory bowel disease and rheumatoid arthritis (Cunninghame Graham et al., 2006; Farh et al., 2015; Kiesler et al., 2009; Leon Rodriguez et al., 2016; Maurano et al., 2012; van Heel et al., 2002; Vince et al., 2016). Other studies implicate polymorphisms at the *Ocab* (*Pou2af1*) locus itself in different forms of autoimmunity (Games Collaborative et al., 2006; Nakamura et al., 2012). We therefore hypothesized that targeting OCA-B would inhibit autoreactive, diabetogenic T cell responses.

Here, we generate a T cell conditional mouse model of OCA-B deficiency. We show that OCA-B ablation protects animals from spontaneous, polyclonal T1D. Protection was associated with reduced autoreactivity of autoantigen-specific CD8^+^ T cells in the pancreatic lymph nodes (PLNs), reduced representation of these same TCR clones in the pancreatic islets and the induction of anergy in potentially autoreactive islet CD4^+^ T cell clones. The protection associated with OCA-B loss varied among different TCR transgenic and monoclonal antigen models, and was weakest using synthetic antigens and monoclonal transplant models. Protection was stronger using a spontaneous autoantigen-specific TCR transgenic model. Using a rational-design approach, we developed membrane-penetrating OCA-B peptide inhibitors that are efficacious in vitro and in mice. Collectively, the results show that OCA-B is a potent and pharmacologically accessible autoimmune regulator.

## Results

### Generation of an *Ocab* conditional allele and NOD backcrossing

We generated a conditional *Ocab* (*Pou2af1*) mouse allele using embryonic stem cells provided by the knockout mouse project (KOMP) repository. Deletion of the *LoxP* sites creates a null allele by removing the first three exons (Fig. S1A). Breeding chimeric mice resulted in germline transmission and a founder mouse (Fig. S1B, lane 2) that was crossed to a FLP deleter mouse strain (FLP^Rosa26^) to remove the reporter cassette and create a floxed conditional allele (Fig. S1A and B, lane 3). Intercrossing these mice generated *fl/fl* homozygotes (Fig. S1B, lane 4) which were crossed to a germline Cre deleter strain (Cre^Rosa26^) to generate germline null *(D)* alleles (lanes 5 and 6). As expected, homozygous *fl/fl* mice produce normal amounts of both p34 and p35 OCA-B protein isoforms (Fig. S1C, lane 6), while no protein of either isoform is present in *Δ/Δ* spleens (lanes 7-8). These results indicate that the *fl* allele is OCA-B sufficient and the Cre-deleted allele is a null. Crossing the *fl* allele onto a CD4-cre driver, which deletes in both CD4 and CD8 compartments, resulted in efficient deletion in splenic T cells (Fig. S1D). This allele therefore represents a robust system in which to study OCA-B function in T cells.

The conditional allele was generated on a C57BL/6 background. To test the role of OCA-B expression in T cells in T1D emergence, we conducted congenic backcrosses to the NOD strain background. This method allows spontaneous diabetes to be produced rapidly by screening and selectively breeding mice with 13 microsatellite and SNP markers associated with autoimmunity (Serreze et al., 1996). *Ocab* is located on mouse chromosome 9 and distant from any of the *Idd* loci. Following these markers with specific primer-pairs (Table S1), we produced backcrossed animals with all 13 markers (Fig. S1E) that recapitulate spontaneous autoimmunity (not shown). For example, *Idd1*, which maps to the MHC region (Hattori et al., 1986), converted to the homozygous NOD allele after 2 backcross generations (Fig. S1E, lane 3), whereas *Idd13*, which maps near beta-2-microglobulin (Serreze et al., 1998) became NOD homozygous after 4 generations (lane 4). Genotyping identified this fourth-generation backcrossed animal as having all 13 microsatellite markers and the *Ocab* conditional allele (Fig. S1F, #122). This founder animal was crossed to NOD.CD4-cre in order to delete OCA-B in T cells. Similar NOD backcrosses, conducted using *Ocab* null mice (Kim et al., 1996), resulted in OCA-B germline-deficient NOD mice (not shown). As with the original C57BL/6 OCA-B germline knockout (Kim et al., 1996), all alleles were viable and fertile with no obvious health issues.

### Normal baseline T cell Status in NOD mice lacking OCA-B

To study the effects of T cell-specific OCA-B loss on T cell function, we performed single-cell RNA sequencing (scRNAseq) and TCR clonotype analysis using CD3^+^ T cells isolated from PLNs of prediabetic 8 wk-old female NOD.*Ocab^fl/fl^*CD4-cre or NOD.*Ocab^fl/fl^* littermate controls. For each group, cells from 3 animals were combined for microfluidics, sequencing and analysis. ~6000 knockout and ~3000 control cells were identified in this analysis, most of which comprise naïve CD4^+^ and CD8^+^ cells (Fig. S2A, pink and orange), and regulatory T cells (Tregs, green). There was also a small population of activated CD4^+^+CD8^+^ cells (blue-green), presumably in the process of exiting the lymph nodes. Other than a small decrease in the percentage of activated T cells (4.4 vs 6.1%), few differences in relative abundance of the populations were observed. None of the populations expressed measurable amounts of *Ocab* (*Pou2af1*) mRNA (not shown), however *Pou2af1* is expressed at low levels in T cells making it difficult capture by scRNAseq. In addition, few changes in gene expression were identified (Fig. S2B, Table S2). We also determined TCR repertoires, identifying few changes across all populations (Fig. S2C). However, specific sub-populations did show changes in TCR utilization. For example, >90% of NOD CD8^+^ T cells that display reactivity to islet-specific glucose-6-phosphatase catalytic subunit related protein (IGRP) residues 206-214 express *Trbv13-3* (Vβ8.1) (Wong et al., 2006). Although this clone was unchanged across the entire population of T cells (Fig. S2C), it was over-represented in the effector CD8^+^ population (Fig. S2D). In contrast, other TCR clonotypes such as *Trbv13-1, Trbv13-2, Trbv1* and *Trbv15* were unchanged (not shown).

### NOD mice lacking OCA-B are protected from T1D

T1D onset in NOD mice is spontaneous and acute (Makino et al., 1985). To test the impact of OCA-B loss in T cells on spontaneous T1D, we compared incidence in in female NOD.*Ocab^fl/fl^*CD4-cre mice, and littermate controls lacking CD4-cre. Glucose was monitored 2-3 times/wk. Approximately 60% of *Ocab^fl/fl^* control mice manifested spontaneous diabetes by 24 weeks of age, reaching an average of >300 mg/dL by 24 weeks, while no *Ocab^fl/fl^*CD4-cre mice became diabetic (Fig. 1A,B). Similar results were observed using female whole-body knockout NOD.*Ocab^-/-^* compared to *Ocab^+/+^* littermate controls. While ~80% of control females developed T1D within 24 weeks after birth, only ~20% of NOD.*Ocab^ΔΔ^*females did so (Fig. 1C). The differences in T1D penetration are likely attributable to differences in the NOD backcrosses, as the 13 *Idd* NOD determinants largely but not completely dictate the autoimmune phenotype. Cumulatively, the results show that OCA-B expression in T cells is critical for T1D pathogenesis in NOD mice.

**Figure 1.**
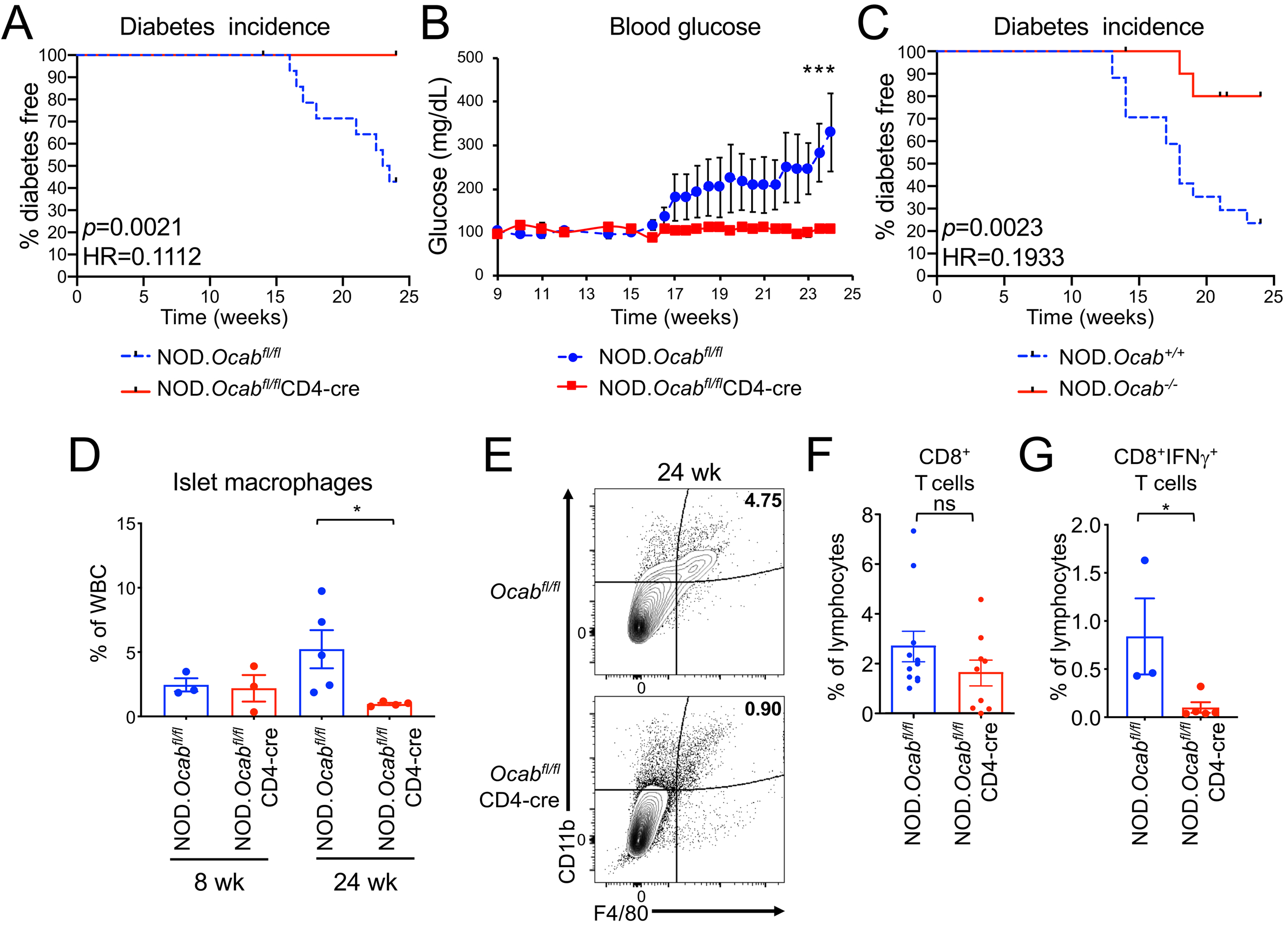
Loss of OCA-B protects NOD mice from T1D. **(A)** Kaplan-Meier plot of diabetes-free survival in littermate female NOD.*Ocab^fl/fl^*CD4-cre (n=12) and NOD. *Ocab^fl/fl^* (n=14) mice. HR=hazard ratio. **(B)** Average blood glucose levels from mice shown in panel A. Student T-test *p*-values: day 23, 0.0458; day 24, 0.0403; day 25, 0.494. **(C)** Percent diabetes-free survival in germline knockout NOD.*Ocab^-/-^* (n=16) and control NOD.*Ocab^+/+^* (n=17) mice was plotted. **(D)** Pancreatic islet leukocytes were isolated from 8 wk- or 24 wk-old littermate female NOD.*Ocab^fl/fl^*CD4-cre (8 wk; n=3, 24 wk; n=5) or NOD.*Ocab^fl/fl^* (8 wk; n=3, 24 wk; n=4) mice and analyzed by flow cytometry. Mean CD45^+^CD11b^+^F4/80^+^ cell frequencies are depicted. Student T-test *p*-value=0.0397. **(E)** Frequencies of CD45^+^CD11b^+^F4/80^+^ cells from representative animals in D are shown. Plots were gated on CD45. **(F)** Mean percentages of total pancreatic-infiltrating CD8^+^ T cell in 9 experimental NOD.*Ocab^fl/fl^*CD4-cre or 11 littermate control NOD.*Ocab^fl/fl^* 12 wk-old mice are plotted. Cells were gated on CD45. **(G)** IFNg expressing CD8^+^ T cell percentages in 16 wk old-NOD.*Ocab^fl/fl^*CD4-cre (n=3) and littermate control islets (n=5) are shown. Student T-test *p*-value=0.0477.

We sacrificed *Ocab^fl/fl^*CD4-cre female mice and controls at 24 weeks, and determined immune cell frequencies in the pancreata. We observed more T cells in the pancreata of OCA-B deficient mice (not shown). An relative increase in infiltrating T cells in endpoint knockouts at endpoint is understandable because the controls have annihilated their pool of beta cells and associated antigens. We also studied pancreatic macrophages in these mice. Islet infiltration of F4/80^+^CD11b^+^ macrophages is critical for NOD T1D pathogenesis both in spontaneous and adoptive transfer T1D models (Peterson et al., 1998; Rosmalen et al., 2000). CCL1 expressed in diabetogenic CD4^+^ T cells recruits CCR8-expressing macrophages (Cantor and Haskins, 2007). OCA-B deletion decreases *Ccl1* mRNA expression in stimulated, rested and re-stimulated CD4^+^ T cells compared to controls (Table S3). Despite the presence of antigen greater numbers of T cells, we observed fewer macrophages in 24 wk NOD.*Ocab^fl/fl^*CD4-cre islets (Fig. 1D,E). In contrast, macrophages were unaffected by T cell-specific OCA-B loss in prediabetic (8 wk) mice (Fig. 1D).

IFNγ is a direct OCA-B target gene (Shakya et al., 2015). Serum and pancreatic IFNγ levels gradually rise in young NOD mice, reaching maximum at diabetes onset (Schloot et al., 2002). Diabetogenic CD4^+^ and CD8^+^ T cells express IFNγ both in humans and NOD mice (Krishnamurthy et al., 2008; Newby et al., 2017; Wang et al., 1997). We therefore predicted that OCA-B loss would reduce IFNγ expression in islet CD4^+^ and CD8^+^ T cells. We compared IFNγ-expressing pancreatic CD4^+^ and CD8^+^ T cells in female NOD.*Ocab^fl/fl^*CD4-cre to littermate control mice lacking CD4-cre. In 12 wk prediabetic mice, CD8^+^ T cell frequencies were only slightly decreased (Fig. 1F). CD4^+^ T cells at the same timepoint were also unchanged (not shown). At the time of diabetes onset (16 wk), fewer IFNγ-expressing CD8^+^ T cells were present in *Ocab* conditional-deleted mice compared to controls (Fig. 1G).

### T cell-specific OCA-B deletion alters the islet-infiltrating TCR repertoire

We performed scRNAseq and TCR clonotype analysis similar to Fig. S2, but using older (12 wk) prediabetic NOD.*Ocab^fl/fl^*CD4-cre or NOD.*Ocab^fl/fl^* littermate control female mice, and using total CD45^+^ cells isolated from pancreatic islets. Cells were isolated from 4 animals/group and combined for microfluidics and sequencing. After filtering, ~6200 knockout and ~3800 control cells were identified. Overlaying cells from both genotypes, we used their gene expression profiles to definitively identify cell lineages, including naïve T cells, memory/effector CD4^+^ T cells, memory/effector CD8^+^ T cells, B cells, macrophages and neutrophils (Fig. 2A). Among the different populations, the biggest relative change was neutrophils, which decreased from 3.4 to 1.4% in conditional knockout islets (Fig. 2A). B cells were also decreased, albeit more slightly. These changes were accompanied by increased percentages of naïve T cells (32 vs. 25%). Prediabetic islets also showed significant changes in gene expression between the 2 groups (Fig. 2B), the strongest of which occurred in neutrophils (e.g., *Cxcl2*, down), memory/effector CD8^+^ T cells (*Ccl4, Tcrg-v1, Itga*, down; *Tcrbv1, Trav8n-2*, up), NK T cells (*Klra2, Lars2*, down) and γδT cells (*Gzma, Gzmb, Ccl5*, down).

**Figure 2.**
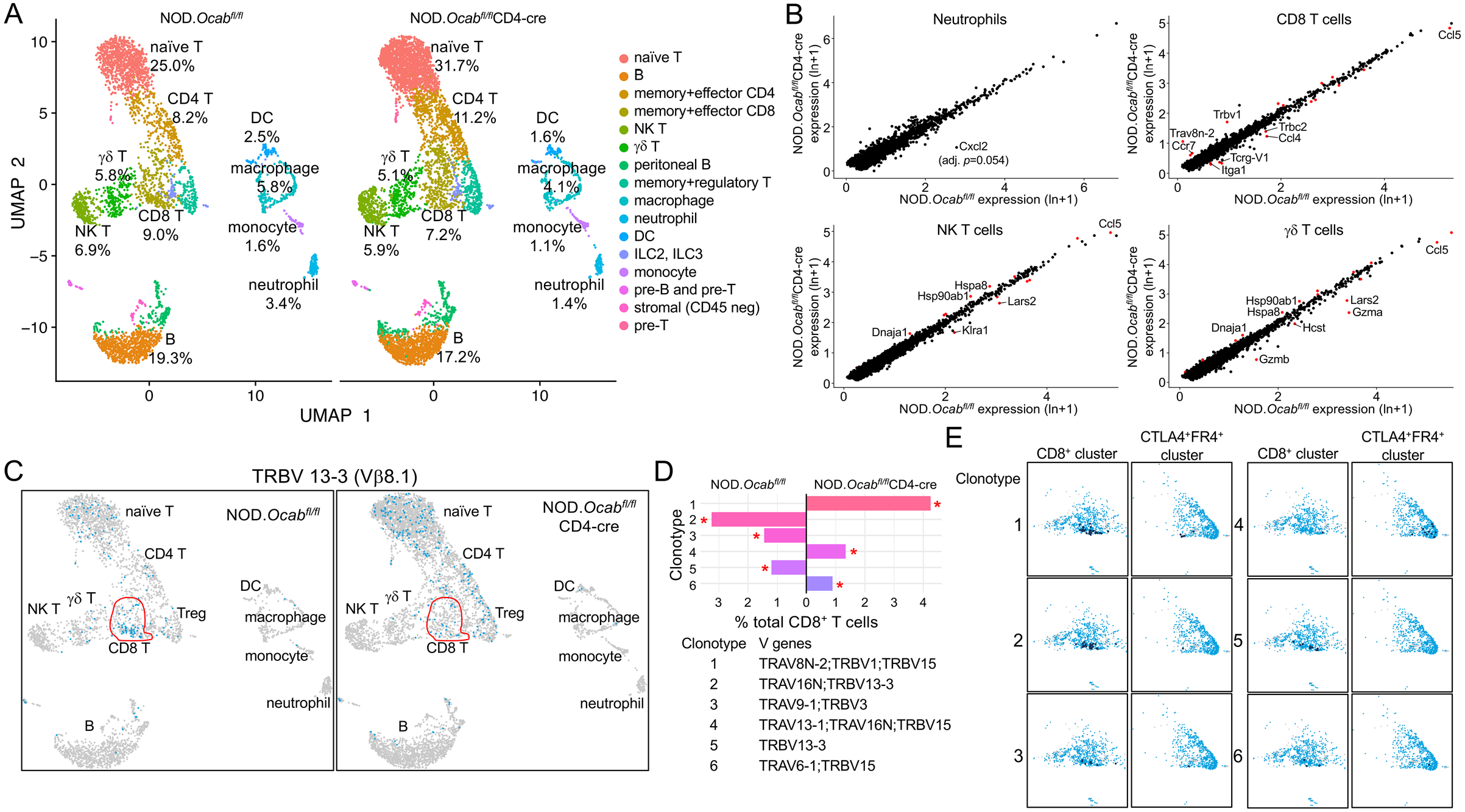
T cell conditional OCA-B loss reduces the numbers of activated, autoreactive pancreatic islet T cells. **(A)** scRNAseq using total islet CD45^+^ cells from prediabetic NOD.*Ocab^fl/fl^*CD4-cre or littermate control NOD.*Ocab^fl/fl^* mice (n=4 for each group). Cell populations were plotted using UMAP (Seurat R package) and percentages in each cluster are shown for each genotype. Clusters were identified using the Seurat R package function FindMarkers. **(B)** 4 clusters from A were analyzed for differential gene expression. Identified genes are shown as a scatter plot. Significantly differentially expressed genes (adjusted *p*-value <0.05) are shown in red. For *Cxcl2* in neutrophils, *p*=3.75×10^-6^, adjusted *p*=0.055. **(C)** UMAP plots similar to A, except cells expressing TCR clonotype 13-3 are shown. The non-naïve (activated+memory) CD8^+^ cell population identified in A is shown in red. **(D)** Percent contribution of the top 6 identified TCR clonotypes to total activated+memory CD8^+^ cells is shown. V genes comprising the clonotypes are shown below. **(E)** Cells positive for clonotypes 1-5 are shown for two clusters, non-naïve (activated+memory) CD8^+^ T cells and CTLA4^+^FR4^+^ anergic cells (consisting of mostly CD4^+^ cells). For each cluster, positive cells are shown in dark blue. An overlay of control and OCA-B deficient cell populations is shown. Each clonotype is only observed in a single genotype (panel D), allowing all cells to be mapped back entirely to OCA-B deficient (clonotypes 1 and 4) or control (clonotypes 2, 3 and 5).

We also profiled TCR utilization in pancreatic T cells. Across total T cells, few changes were observed (Fig. S3), however this was largely do to similar representation in the naïve T cell pool. Isolating antigen-experienced CD4^+^ and CD8^+^ effector + memory subpopulations revealed sharp differences. For example, >90% of diabetogenic CD8^+^ T cells displaying reactivity to IGRP206-214 express *Trbv13-3* (Vβ8.1) (Wong et al., 2006). Relative to controls, this clonotype was under-represented in OCA-B deficient effector + memory CD8^+^ cells (Fig. 2C, 20.9% vs 8.6%). Based on this information, we repeated the clonotype analysis, but only using the effector + memory CD8^+^ T cell cluster. Within this population, dominant TCR clonotypes were strikingly different (Fig. 2D). Clonotype 1 for example is the most abundant, and corresponds to a TCRa V8N-2 chain paired with TCRβ V1 or V15. It has not been associated with T1D. This clonotype was absent in control islets but represented >4% of OCA-B deficient islet CD8^+^ effector + memory T cells (Fig. 2D). Clonotype 1 cells overlaid with both genotypes is shown in Fig. 2E. Clonotype 2 by contrast expresses the T1D-associated Vβ8.1 TCR β chain. This clonotype was strongly represented in controls (>3%) but absent from OCA-B deficient effector + memory CD8^+^ cells (Fig. 2D). In contrast to CD8, CD4-dominant TCR clonotypes associated with T1D were still present in knockouts, but associated with markers of anergy. For example, clonotype 4 (*Trav13-1/16N+Trbv15*) is associated with pro-diabetogenic, BDC-reactive CD4^+^ T cells (Li et al., 2009). This clonotype was enriched in the OCA-B deficient effector + memory CD4^+^ cells, but only in a CTLA4- and FR4-expressing sub-population (Fig. 2E). Cumulatively, these results show that OCA-B loss from T cells significantly skews the TCR landscape of islet effector/memory-like T cells to an anti-diabetogenic phenotype.

### Autoantigen-specific CD8+ T cells are present in PLNs of OCA-B deficient mice, but show reduced reactivity and islet penetration

In order to understand the basis for the reduction in autoantigen-specific CD8^+^ T cell islet clones, we studied Vβ8.1 expression and IGRP_206-214_ reactivity in matched pancreatic islets and PLNs from 12 wk prediabetic female *Ocab^fl/fl^*CD4-cre mice or littermate *Ocab^fl/fl^* controls. Islet CD8^+^ T cells showed decreased Vβ8.1 usage (Fig. 3A), while PLNs from the same animals showed a corresponding increase (Fig. 3B). To directly test diabetogenic, antigen-specific islet CD8^+^ T cells, we used IGRP_206-214_ H-2K^d^ class I tetramers, which recognize TCRs expressing Vβ8.1. Islet CD8^+^Vβ8.1^+^ T cells showed a strong decrease in IGRP_206-214_ specificity (Fig. 3C).

**Figure 3.**
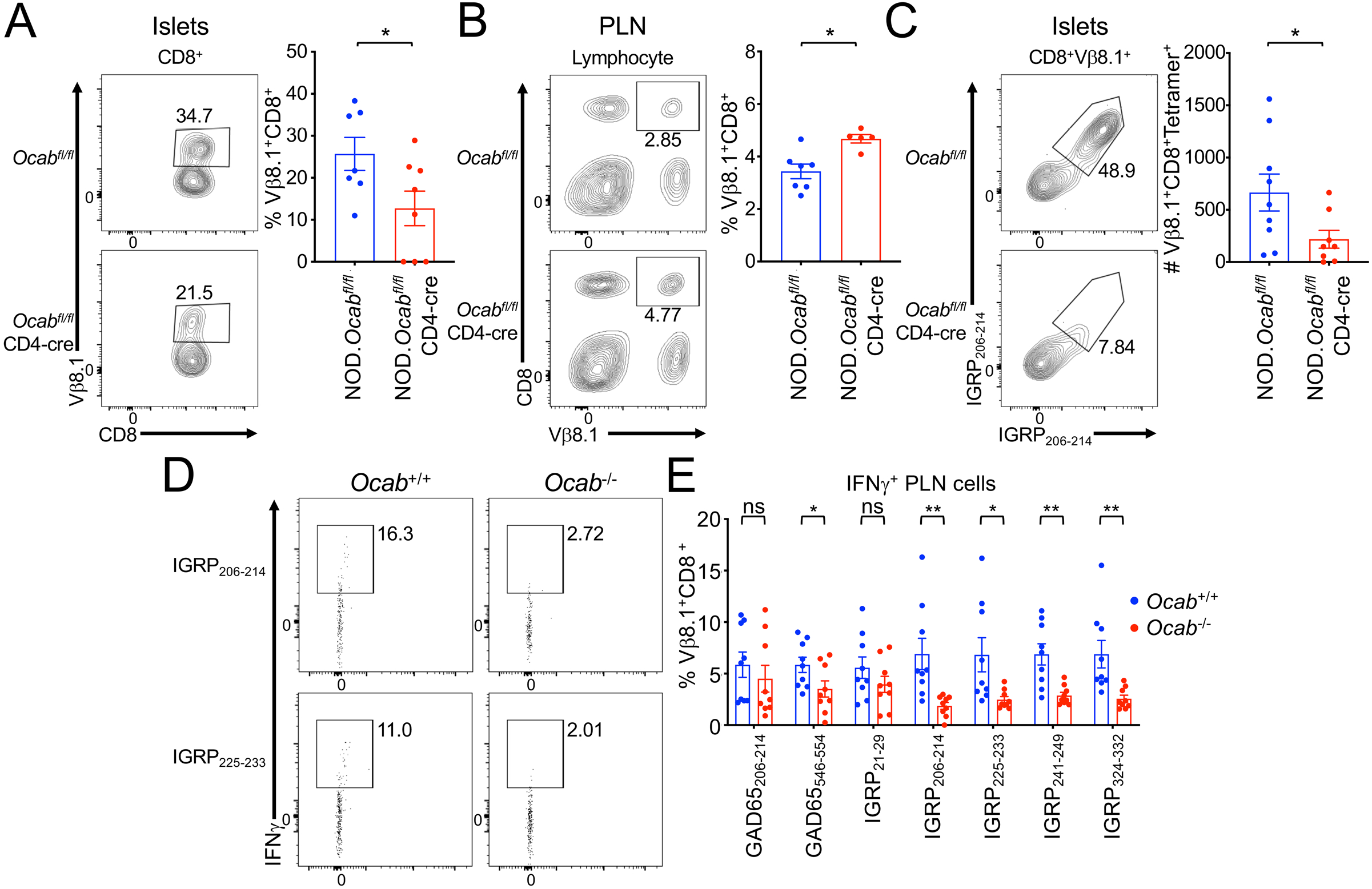
Autoantigen-specific CD8+ cells become nonreactive and fail to infiltrate islets in the OCA-B deficient condition. **(A)** Pancreatic leukocytes were isolated from prediabetic 12 wk littermate female NOD.*Ocab^fl/fl^*CD4-cre (n=9) or NOD.*Ocab^fl/fl^* (n=10) mice and analyzed by flow cytometry. Representative plots (left) and total CD8^+^Vβ8.1^+^ T cell numbers (right) are shown. Student T-test *p*-value=0.030. **(B)** PLN leukocytes were isolated from same mice (NOD.*Ocab^fl/fl^*CD4-cre, n=7 or NOD.*Ocab_fl/fl_* n=5) in A. Representative plots (left) and total CD8^+^Vβ8.1^+^ T cell frequencies (right) are shown. Student T-test *p*-value=0.030. **(C)** Representative plots (left) and total cell numbers (right) of CD8^+^Vβ8.1^+^ H-2K^d^ IGRP_206-214_ tetramer-positive islet T cells are depicted from NOD.*Ocab^fl/fl^*CD4-cre (n=9) or NOD.*Ocab^fl/fl^* (n=10). Student T-test *p*-value=0.040. **(D)** Total PLN WBCs were isolated from prediabetic 12 wk-old littermate NOD.*Ocab^-/-^* or NOD.*Ocab^+/+^* mice and re-stimulated with IGRP peptides (IGRP_206-214_ or IGRP225-233). Cells were analyzed for IFNγ expression by flow cytometry. **(E)** Mean percentages of cells expressing IFNγ from 9 experiments (3 replicates from 3 *Ocab^+/+^* and 3 *Ocab^-/-^* mice) for the peptides in D, as well as two GAD65 peptides (206-214 and 546-554) and three additional IGRP peptides (21-29, 241-249 and 324-332). Significant student T-test *p*-values were: GAD65_546-554_, 0.046; IGRP_206-214_, 0.005; IGRP_225-233_, 0.030; IGRP_241-249_, 0.004; IGRP_324-332_, 0.012.

The decrease in autoantigen-specific islet CD8^+^ T cells and corresponding increase in the PLNs in OCA-B T cell deficient mice raised two possibilities: cells in the PLNs could fail to become activated, or could become activated but fail to migrate to the pancreas. In the first case, more antigen-reactive cells would be observed while in the second there would be fewer. To distinguish between these possibilities, we performed peptide stimulation experiments using 7 different known diabetogenic self-peptides. We used 5 IGRP peptides, including IGRP_206-214_ which corresponds to the tetramer used above and the NY8.3 TCR transgenic mice used below. We also used two GAD65 epitopes, 206-214 and 546-554. 5 of the 7 peptides showed significant reductions in autoreactivity, including IGRP_206-214_ (Fig. 3D,E). These results indicate that potentially autoreactive cells fail to become activated in the PLNs of OCA-B deficient mice.

### Protection conferred by OCA-B loss is model-specific

T1D in NOD mice originates from defective negative selection in developing T cells, resulting in T cell autoreactivity. OCA-B is not expressed in thymocytes and OCA-B loss does not appear to affect T cell development (Shakya et al., 2015). Therefore, the protective effect of OCA-B loss in T cells likely arises from blunting pre-existing autoreactivity in the periphery. To test the hypothesis that OCA-B loss confers protection in simplified systems, we crossed OCA-B deficient mouse models employing TCR transgenes and model antigens.

We crossed the germline null *Ocab* allele to BDC2.5 mice (Katz et al., 1993), which expresses a CD4-restricted TCR specific for a hybrid insulin-chromogranin A peptide (Delong et al., 2016). Transferring Treg-depleted T cells from BDC2.5 TCR transgenic mice into NOD.Scid mice results in disease after approximately 10 d (Berry and Waldner, 2013; Presa et al., 2015). We transferred CD4^+^CD25^-^ T cells from germline *Ocab^-/-^* or *Ocab^+ +^* BDC2.5 mice into NOD.Scid recipients, monitoring diabetes emergence. Transplants from both *Ocab^-/-^* and control NOD.BDC2.5 mice resulted in equivalent disease kinetics (Fig. 4A) and severity (not shown). Transferring *Ocab^-/-^* or control NOD.BDC2.5 CD4^+^CD62L^+^Vβ4^+^ T cells into NOD.Scid recipients, or total CD4^+^ T cells into NCG recipients, also resulted in equivalent disease (Fig. S4A, Fig. 4B).

**Figure 4.**
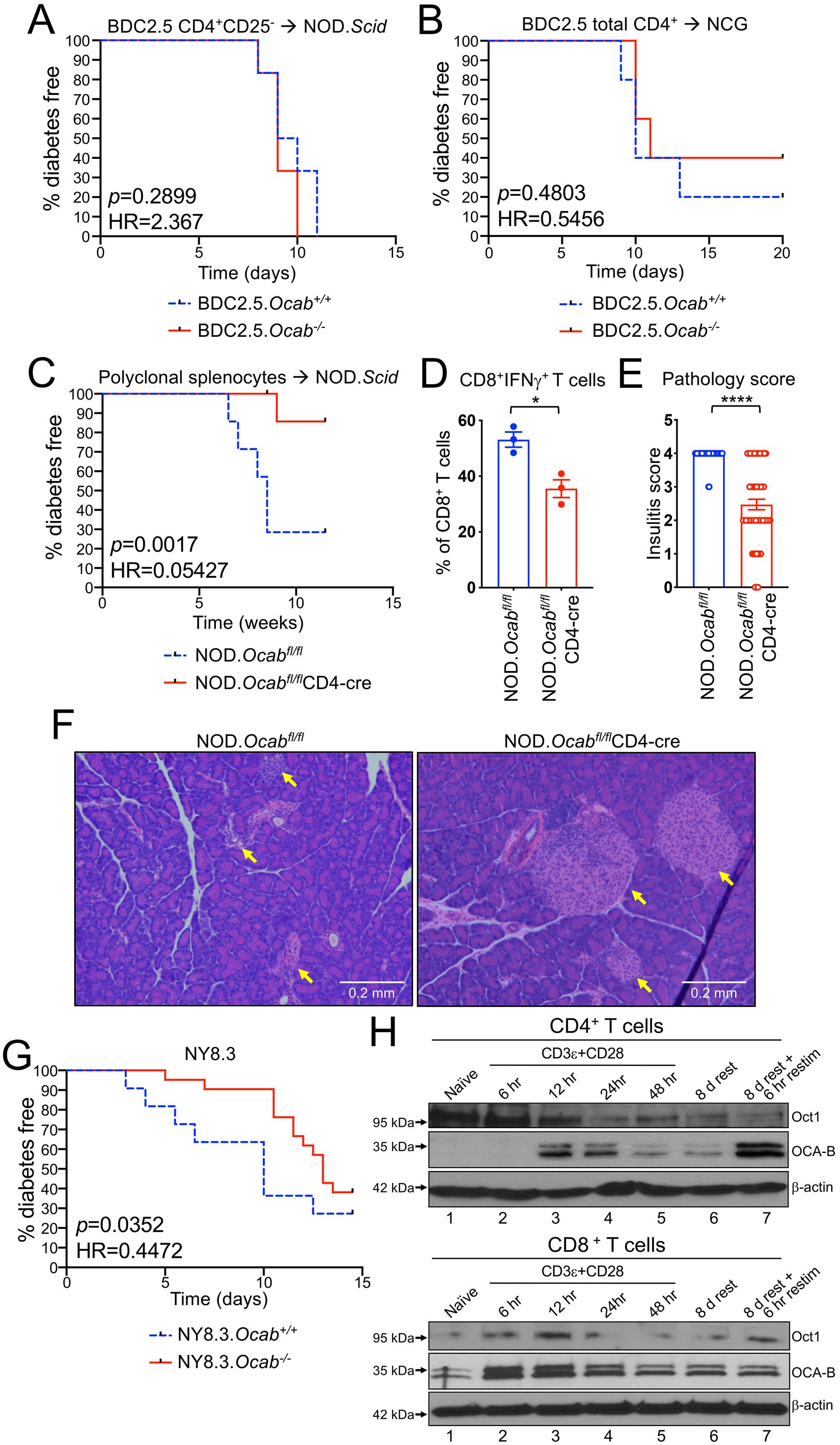
OCA-B loss protects NOD mice from T1D in spontaneous and polyclonal splenocyte transfer models but not monoclonal transfer models. **(A)** 2×10^5^ purified CD4^+^CD25^-^ splenic T cells from NOD.BDC2.5.*Ocab^-/-^* or control NOD.BDC2.5.*Ocab^+/+^* donors were injected retro-orbitally into NOD.Scid (n=6 for each group) mice. Diabetes-free survival is shown. **(B)** 1.5×10^6^ purified splenic total CD4^+^ T cells from NOD.BDC2.5.*Ocab^-/-^* or control donors were transferred into NCG mice (n=5 for each group). Mice were monitored for diabetes development. **(C)** Total NOD splenocytes (5×10^6^) from prediabetic 6-to 8-wk-old NOD.*Ocab^fl/fl^*CD4-cre or control NOD.*Ocab^fl/fl^* donors were adoptively transferred into sex-matched NOD.Scid (n=7 for each group) recipients. T1D-free survival is shown. **(D)** 13 wk post-transfer, the proportion of IFNγ-expressing islet CD8^+^ T cells was assessed by flow cytometry from mice in C. Student T-test *p*-value=0.0136. **(E)** Pancreata from the same mice as in C were fixed, sectioned and H&E stained. Pathological scores are shown based on 6-7 islets/slide, 3 slides/mouse and 3 mice/group (>60 islets/group). Student T-test *p*-value=1.04×10^-5^. **(F)** Example pancreatic images from endpoint animals. Yellow arrows indicate islet positions. Images were collected at 10× magnification. **(G)** The *Ocab* null allele was crossed to NY8.3 TCR transgenic mice. Spontaneous T1D was measured in female *Ocab^-/-^* (n=20) or control *Ocab^+/+^* (n=12) littermates. **(H)** Naïve CD4^+^ or CD8^+^ T cells (CD8a^neg^ or CD4^neg^, CD11b^neg^, CD45R^neg^, DX5^neg^, Ter-119^neg^, CD44^lo^) were isolated from C57BL/6 spleens and stimulated for up to 2 d in vitro using anti-CD3ε and CD28 antibodies. Cells were then washed and replated in the presence of exogenous IL-2. After 8 d rest in culture, cells were restimulated for 6 hr. Lysates were prepared from each step and subjected to OCA-B immunoblotting to assess changes in expression. Oct1 and β-actin are shown as controls.

We also tested the effect of T cell conditional OCA-B deletion in a different strain background, C57BL/6J, using the artificial membranous chicken ovalbumin antigen expressed from the rat insulin promoter (RIP-mOVA) (Van Belle et al., 2009). Wild-type OVA-reactive CD8^+^ OT-I T cells were transferred into either *Ocab^fl/fl^*CD4-cre or control *Ocab^fl/fl^* RIP-mOVA mice. In this model, host CD4^+^ T cells help transferred OT-I T cells promote T1D pathogenesis (Kurts et al., 1997). Mice were infected with Lm-OVA to induce strong and synchronous beta cell destruction and loss of glucose control (>400 mg/dL). No differences were observed using host mice with or without OCA-B T cell deletion (Fig. S4B). These results indicate that OCA-B does not act analogously to transcription factors that regulate many of the same genes, such as NF-AT. Knockouts of these factors would result in broad immunosuppression and protection from T1D across models. Rather, our results suggest that OCA-B loss selectively blunts immune activation in contexts such as autoantigen recognition. To test this supposition and rule out the possibility that T cell transfer itself does not blunt the effect of OCA-B loss, we transferred total splenocytes from polyclonal NOD.*Ocab^fl/fl^*CD4-cre mice or controls. This model, which takes weeks rather than days to develop T1D, recapitulates robust T1D protection with T cell-specific OCA-B loss (Fig. 4C). Protection was associated with fewer IFNγ-expressing islet cytotoxic T cells (Fig. 4D), and reduced insulitis in pancreatic histological sections (Fig. 4E,F).

Because the Vβ8.1 clonotype and IGRP_206-214_-reactive cells were significantly attenuated in the islets of T cell conditional OCA-B deficient mice (Fig. 2C-E, Fig. 3A-C), we crossed the *Ocab* germline allele to the NY8.3 TCR transgenic line, which expresses a CD8-restricted TCR directed towards IGRP_206-214_ and manifests spontaneous T1D (Verdaguer et al., 1997). We passively monitored *Ocab^-/-^* or control *Ocab^+/+^* NY8.3 mice for T1D development. Significant protection was observed (Fig. 4G, HR=0.4472, *p* = 0.0352), though the degree of protection was less than in OCA-B germline knockout polyclonal models. Together with the decreased IGRP reactivity in OCA-B deficient PLNs, this result indicates that OCA-B promotes T1D in peripheral auto-reactive CD8^+^ T cells. Using immunoblotting with OCA-B antibodies, we confirmed that OCA-B is expressed in primary splenic C57BL/6 CD8^+^ T cells. Unlike CD4^+^ cells where OCA-B is undetectable in naïve cells but becomes induced with prolonged activation (Fig. 4H, top panels), OCA-B^+^ could be detected in naïve CD8^+^ cells was rapidly augmented upon activation (Fig. 4H, lanes 1-2). Similar to CD4^+^ cells, expression was stably maintained when the stimulus was removed and cells were rested with IL-2 (lane 6).

### OCA-B deficiency promotes CD4^+^ T cell anergy in vitro

CD8-restricted TCR clones associated with T1D are depleted from the islets of OCA-B T cell conditional mice. In contrast, CD4-restricted clones are increased but associated with an anergic phenotype (Fig. 2E). Anergy is a peripheral tolerance mechanism in which T cells that encounter antigen without co-stimulation become nonfunctional (Kalekar et al., 2016; Kearney et al., 1994; Vanasek et al., 2001). Normal naïve primary CD4^+^ T cells can be activated in culture by TCR and costimulatory receptor activation, e.g. with anti-CD3ε/CD28 antibodies. Providing TCR signals without CD28 antibodies generates anergic responses (Chai and Lechler, 1997). Interestingly, OCA-B is induced to the same extent with or without co-stimulation (Fig. 5A). To determine if OCA-B loss promotes anergic CD4^+^ T cell responses, we stimulated naïve splenic CD4^+^ T cells from germline knockout *Ocab^-/-^* and control *Ocab^+/+^* animals ex vivo using plate-bound anti-CD3ε antibodies without co-stimulation, and restimulated the cells with PMA/ionomycin. IL-2 responses were weak in restimulated cells, as expected given the absence of initial costimulation. CD4^+^ T cells lacking OCA-B generated ~2.5-fold less IL-2 (Fig. 5B,C), indicating that OCA-B provides a barrier against anergic responses.

**Figure 5.**
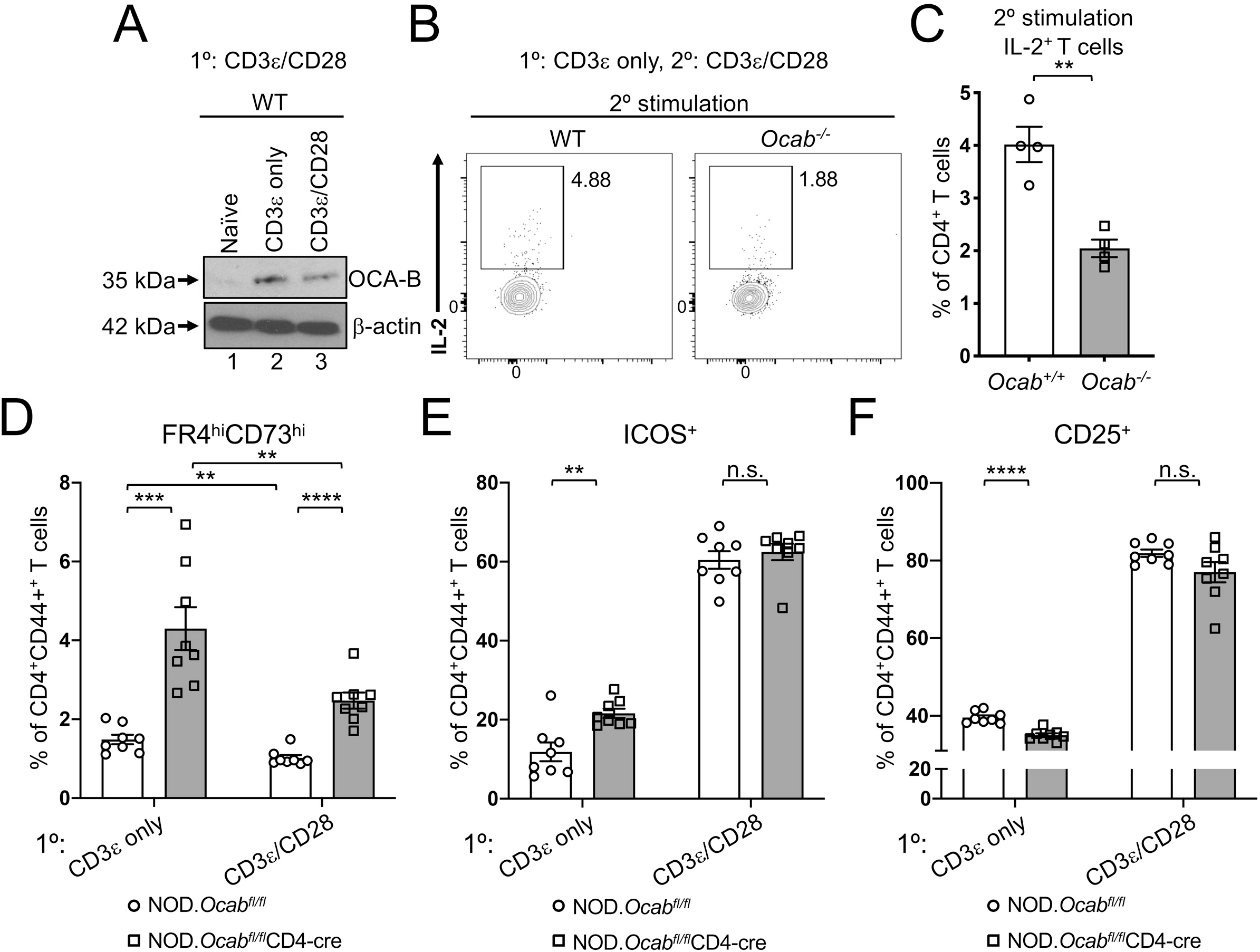
OCA-B loss in CD4^+^ T cells increases anergy in vitro. **(A)** Naïve CD4^+^ C57BL/6 T cells were stimulated in vitro for 24 hr with anti-CD3ε antibodies ± co-stimulation with CD28 antibodies. Lysates were prepared and subjected to immunoblotting using OCA-B antibodies. β-actin is shown as a loading control. **(B)** Naïve OCA-B deficient and control CD4^+^ T cells were stimulated in vitro with anti-CD3ε antibodies. 48 hr later, the cells were re-stimulated with PMA and ionomycin for 6 hr in the presence of brefeldin A, stained for intracellular IL-2 and analyzed by flow cytometry. Cells were gated on CD4 and CD44. **(C)** Quantitation using independently purified cells from the spleens of 4 mice treated similar to B. Student T-test *p*-value=0.0019. **(D)** Naïve OCA-B deficient and littermate control splenic CD4^+^ T cells from 6 wk-old NOD mice were stimulated for 2 d in vitro with indicated antibodies, replated and rested in the absence of antibody for 2 d, and analyzed by flow cytometry. Mean FR4^hi^ CD73^hi^CD4^+^CD44^+^ cell frequencies are shown using independently purified cells from the spleens of 2 mice, with 4 independent culture replicates performed for each mouse (n=8). Student T-test *p*-values: CD3ε only, 0.0002; CD3ε/CD28, 1.04×10^-5^; control CD3ε vs. CD3ε/CD28, 0.0053; OCA-B deficient CD3ε vs. CD3ε/CD28, 0.0068. **(E)** Similar to D, except frequencies of CD4^+^CD44^+^ICOS^+^ cells are plotted. CD3ε only student T-test *p*-value=0.0026. **(F)** Similar to D, except average percentage of CD4^+^CD44^+^CD25^+^ cells are plotted. CD3ε only student T-test *p*-value=1.80×10^-5^.

In CD4^+^ cells, ICOS (Inducible T-cell costimulator) promotes the induction of T cell anergy (Dong et al., 2001; Dong and Nurieva, 2003; Tuettenberg et al., 2009) and is up-regulated by Oct1 loss in naïve T cells subjected to stimulation under anergic conditions (Kim et al., 2019). Oct1 loss also elevates expression of FR4 and CD73 (Kim et al., 2019) the co-expression of which is a marker for anergy (Kalekar et al., 2016; Martinez et al., 2012). To determine whether OCA-B loss similarly affects expression of these proteins, we stimulated naïve splenic CD4^+^ T cells (CD8a^neg^CD11b^neg^CD45R^neg^DX5^neg^Ter-119^neg^CD44^lo^) from 6 wk-old *Ocab^fl/fl^*CD4-cre and littermate control CD4-cre male NOD mice for 2 d ex vivo using anti-CD3ε antibodies ± costimulation. Cells were then rested for an additional 2 d. Across all conditions, CD4^+^CD44^+^FR4^hi^CD73^hi^ anergic cells were present at >2-fold higher frequency in the OCA-B deficient condition (Fig. 5D). The fraction of CD44^+^ cells that express ICOS was similarly elevated in cells lacking OCA-B, though only without costimulation (Fig. 5E). Additionally, the expression of the activation marker CD25 was slightly decreased (Fig. 5F). Cumulatively these results suggest that OCA-B loss in CD4^+^ T cells institutes higher immune thresholds by augmenting anergic responses.

### Generation of an OCA-B peptide inhibitor

The normal T cell developmental and primary immune response phenotypes observed with OCA-B genetic deletion suggested the possibility of a “therapeutic window” in which targeting OCA-B pharmacologically would blunt autoimmunity while minimally affecting baseline immune function. Based on available OCA-B structural information, we generated a membrane-permeable peptide inhibitor of OCA-B’s downstream effector functions. Oct1 interacts with Jmjd1a in a MEK/ERK-dependent manner (Shakya et al., 2015; Shakya et al., 2011) and contains two potential ERK phospho-acceptor serines. The structure of the Oct1 DNA binding domain complexed with consensus octamer binding site DNA (Klemm et al., 1994) reveals that these Oct1 serines are located in the flexible, solvent-exposed linker domain, which connects the two DNA binding sub-domains (Fig. 6A, red dashed line). In contrast with Oct1, OCA-B constitutively interacts with Jmjd1a (Shakya et al., 2015). The OCA-B N-terminus has been solved in complex with the Oct1 DNA binding domain and consensus octamer DNA (Chasman et al., 1999). This region of OCA-B is critical for both Oct1 binding and transcription activity (Gstaiger et al., 1996). To identify potential Jmjd1a-interacting OCA-B regions, we aligned the full-length Oct1 and OCA-B amino acid sequences to a cofactor interaction surface of androgen receptor (AR), which also interacts with Jmjd1a (Yamane et al., 2006). Human AR mutations that cause androgen insensitivity have been mapped to residues 698-721 (Thin et al., 2002). Aligning this span to Oct1 identifies the linker domain as the top hit (Fig. 6B). The alignment shows conservation of 3 out of 4 deleterious AR mutations (asterisks). The mutation site that is not conserved is a potential ERK target serine in Oct1, and a phospho-mimetic glutamic acid residue in AR (blue box). These findings suggest that the Oct1 linker domain constitutes a surface which when phosphorylated interacts with Jmjd1a.

**Figure 6.**
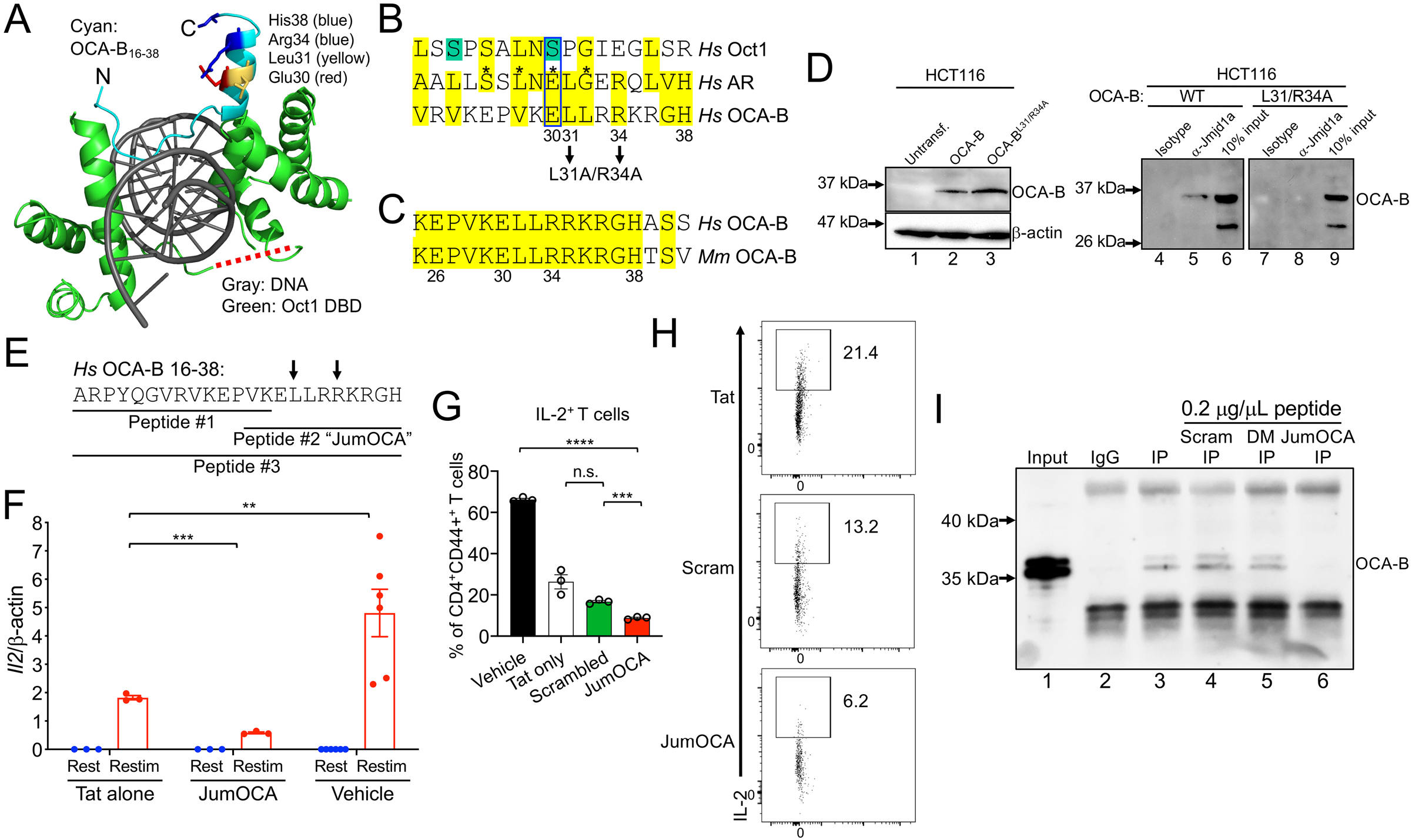
Design and validation of OCA-B peptide inhibitors. **(A)** OCA-B N-terminus (residues 16 and 38)/Oct1 DNA binding domain/octamer binding DNA co-crystal structure (PDB ID 1CQT) (Green, 2016). Gray: DNA. Green: Oct1 DNA binding domain. Cyan: OCA-B. Red dashed line shows position of the Oct1 linker domain. **(B)** Top alignment of the human androgen receptor (AR) isoform transcript variant 1 (Sequence ID: ADD26780.1) co-activator interaction domain (residues 698-721) with full-length human Oct1 and OCA-B. The aligned regions are from the Oct1 linker domain and OCA-B N-terminus. Green serines: putative ERK phospho-acceptor sites. Yellow: similar or identical amino acids. Asterisks: known human point mutations that block coactivator binding and cause androgen insensitivity syndrome in humans. Pairwise alignments were performed using the FASTA algorithm (https://embnet.vital-it.ch/software/LALIGN_form.html) and trimmed for 3-way overlap. **(C)** Alignment of human and mouse primary OCA-B peptide sequences. **(D)** Co-IP of Jmjd1a with L31A/R34A double point-mutant OCA-B and wild-type control. HCT116 cells transfected with WT or mutant OCA-B constructs were used. Protein expression was checked 48 hrs after transfection by Western blotting. 50 μg input protein was loaded in lanes 1-3. **(E)** Indicated peptide sequences were synthesized as C-terminal Tat fusions for membrane permeability. Arrows indicate position of mutant in B and D. **(F)** *Il2* mRNA expression in primary naïve CD4^+^ T cells treated with 50 μM JumOCA peptide was measured relative to β-actin internal standard by RT-qPCR. Cells were stimulated with CD3ε/CD28 antibodies for 2 d, rested for a further 8 d in the presence of exogenous recombinant IL-2, and restimulated for 6 hr. Peptide was included during resting and restimulation only, and replaced every other day with media changes. Three independent biological replicates were used for each condition. Tat vs. JumOCA restim student T-test *p*-value=0.0007. Tat vs. vehicle student T-test *p*-value=0.0022. **(G)** IL-2 cytokine expression in primary naïve CD4^+^ T cells cultured with 50μM peptide at initial treatment, and with 25 μM peptide from the secondary treatment was measured by flow cytometry. Cells were treated similarly to F except collection, brefeldin A treatment and processing for flow cytometry occurred 24 hr post-restimulation. Three independent biological replicates were used for each condition. Vehicle vs. JumOCA student T-test *p*-value=9.62×10^-7^. Scrambled peptide vs. JumOCA student T-test *p*-value=0.0006. **(H)** CD4^+^CD44^+^IL2^+^ T cell frequencies from representative samples in G. Plots were gated on CD4 and CD44. **(I)** Peptide effects on the interaction between OCA-B and Jmjd1a was measured by co-immunoprecipitation. M12 B cells were used. After incubation with protein-G beads, 0.2 μg/μL control scrambled (Scram), double point mutant (DM) peptide or JumOCA peptide were added for a further 3 hr prior to precipitation and washing. 1% input (lane 1) is shown as a control.

As with Oct1, alignment with OCA-B identifies a potential Jmjd1a interacting surface (Fig. 6B). Unlike Oct1, a glutamic acid residue (Glu30) aligns with AR. Furthermore, the OCA-B residues aligning to AR lie on one side of an alpha-helix (Fig. 6A). Glu30 (red), Leu31 (yellow), Arg34 (blue) and His38 (blue) may therefore constitute a Jmjd1a docking surface. These residues are conserved between mouse and human (Fig. 6C). We mutated OCA-B L31 and R34 to alanine in a transient expression plasmid, and transfected the double-point mutant or parent plasmid control into HCT116 cells, which do not express endogenous OCA-B. Control co-IP using Jmjd1a antibodies and lysates from cells expressing wild-type OCA-B confirms the interaction (Fig. 6D, lane 5), while the mutant OCA-B protein failed to interact (lane 8). The mutant was expressed equivalently to WT (lanes 2, 3). These results show the importance of OCA-B residues 30-38 for cofactor interactions.

We synthesized three overlapping peptides corresponding to the OCA-B N-terminus (Fig. 6E, peptides #1, #2 (hereafter called “JumOCA”) and #3), as C-terminal fusions to the HIV Tat protein for membrane permeability (Schwarze et al., 2000). We also conjugated HIV Tat to FITC to enable peptide tracking. Incubating total splenic or pancreatic CD3^+^ T cells in ex vivo culture with 45 μM Tat-fused peptide for 15 min significantly concentrated the peptide within cells. Control OVA-fused peptide lacking membrane-penetrating properties showed no effect (Fig. S5). We then treated splenic CD4^+^ T cells with the Tat-fused JumOCA peptides. Prior work has shown that 6 hr restimulation of resting but previously activated OCA-B deficient CD4^+^ T cells results in decreased expression of target genes such as *Il2* (Shakya et al., 2015). Using this assay with 50 μM JumOCA peptide (every other day with media changes during rest and restimulation) inhibited *Il2* mRNA expression relative to β-actin by ~10-fold (Fig. 6F, compare Tat peptide to JumOCA). Although treated cells showed no obvious changes in viability, morphology or expansion during the course of the assay (not shown), there was a 2-3-fold nonspecific diminution of activity associated with unfused Tat peptide (Fig. 6F, compare Tat peptide with no peptide), which has been associated with toxicity (Caron et al., 2004; Krautwald et al., 2004; Polo et al., 2004). Similar results were obtained with the larger peptide #3 but not with peptide #1 (not shown). Peptide #1 is missing residues important for the Jmjd1a interaction based on mutagenesis (Fig. 6B). For all further experiments, we used the JumOCA peptide.

We then synthesized a Tat-conjugated peptide using a scrambled JumOCA sequence. This peptide has the same amino acid composition, mass and pI, but should not be able to efficiently compete for Jmjd1a. We used the Tat-only, scrambled and JumOCA peptides with flow cytometry to assess IL-2 production in restimulated CD4^+^ T cells. JumOCA significantly inhibited IL-2 production relative to the scrambled control, however there were significant toxicities associated with both the scrambled and Tat-only peptides relative to vehicle (Fig. 6G). An example mouse from this experiment is shown in Fig. 6H.

The above results are consistent with the JumOCA peptide operating via Jmjd1a competitive inhibition. To test this directly, we used scrambled, double point mutant and wild-type (JumOCA) peptides (lacking the Tat membrane penetrating domain) in a Jmjd1a co-IP assay. We subjected lysates from mouse M12 B cells, which expresses endogenous OCA-B, to immunoprecipitation with Jmjd1a antibodies, resulting in OCA-B co-IP (Fig. 6I, lane 3). Incubation of precipitated material with scrambled or double point mutant peptides had no effect on OCA-B co-immunoprecipitation (lanes 45), while the same concentration of JumOCA peptide efficiently blocked OCA-B recovery (lane 6).

### JumOCA protects NOD mice from newly-arisen T1D

By the time symptoms arise in NOD mice, 90-95% of beta cells have been destroyed. Remaining beta cells are rendered nonfunctional due to insulitis. Alleviating this inflammation provides a “honeymoon period” during which glucose homeostasis can be restored. We treated 12-18 wk littermate female NOD mice whose glucose levels were newly-risen above 225 mg/dL but still below 275 mg/dL with three intravenous injections of 10 mg/kg JumOCA peptide or Tat-only peptide control. Injections were spaced 12 hr apart. Immediately after the second injection and 4 hr after the final injection, blood glucose was measured. Strikingly, JumOCA but not Tat-only or scrambled peptide reversed elevated blood glucose (Fig. 7A). Islet T cell pro-inflammatory IFNγ and IL-17A cytokine production were reduced by JumOCA peptide treatment compared to Tat-only controls (Fig. 7B,C). In contrast to the pancreas, PLNs showed no change in T cell numbers or percentages (Fig. 7D) but similar decreases in pro-inflammatory cytokine production (not shown).

**Figure 7.**
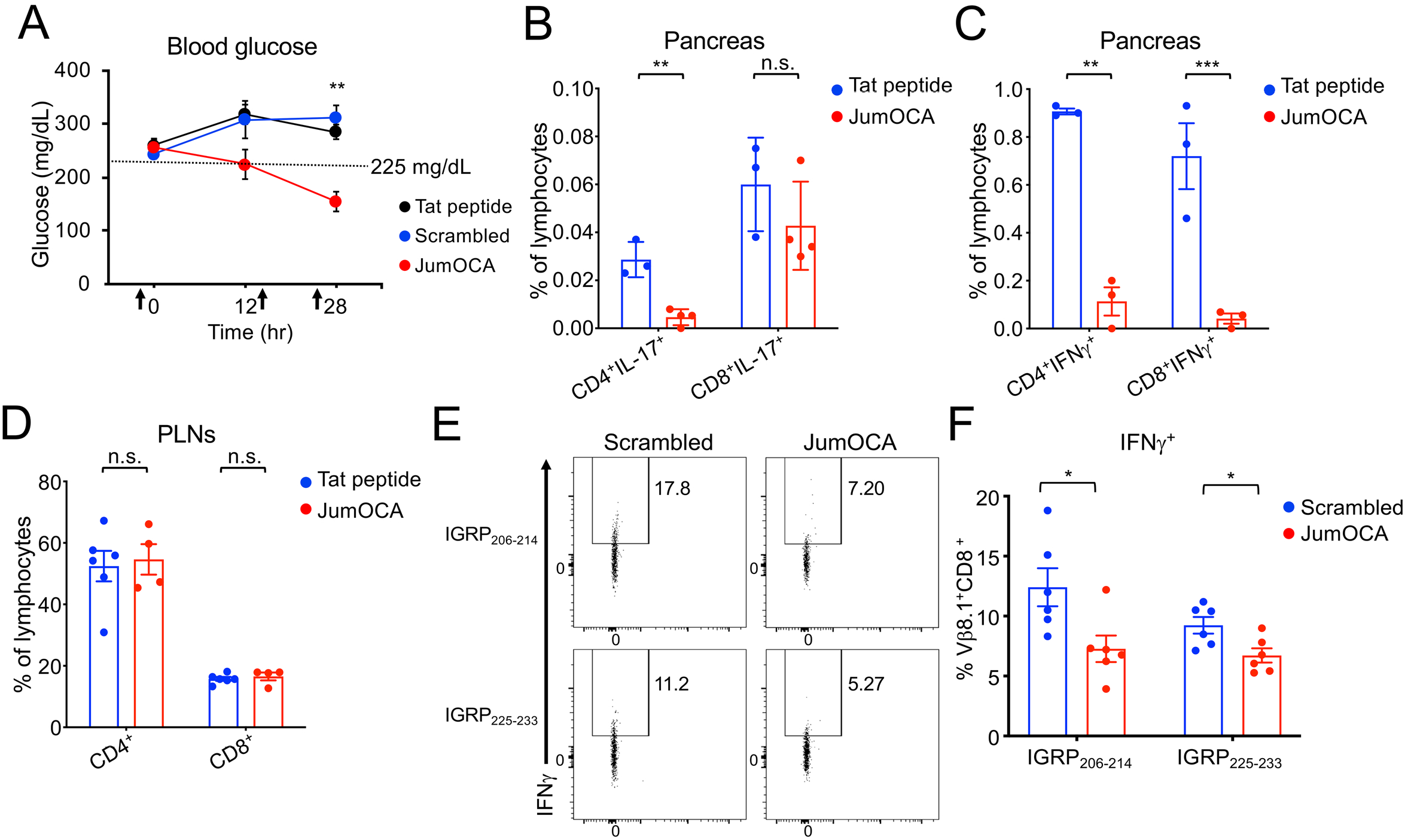
OCA-B inhibitor peptide reduces blood glucose and inflammatory cytokine levels in the pancreas in NOD mice with newly-arisen diabetes. **(A)** 3 doses of 10 mg/kg JumOCA (n=4), Tat only (n=6) or scrambled peptide (n=3) were injected retro-orbitally every 12 hr into 12-18 wk-old NOD female littermate mice whose glucose levels were newly-risen above 225 mg/dL but still below 275 mg/dL. Glucose levels are shown immediately before the first injection, after the second injection and 4 hr after the final injection. Data were collected on a rolling basis as mice became spontaneously diabetic. JumOCA vs. Tat peptide student T-test *p*-value=0.0004. JumOCA vs. scrambled peptide student T-test *p*-value=0.0064. **(B)** 12 hr after the last injection of peptides in A, PLN CD4^+^ and CD8^+^ T cell percentages were analyzed by flow cytometry. Mean of islet IL-17 expressing T cells were analyzed by flow cytometry and plotted. Results are from 3 independent mice in the case of Tat and 4 independent mice in the case of JumOCA. CD4^+^IL-17^+^ student T-test *p*-value=0.002. **(C)** Similar analysis as B except for IFNγ. CD4^+^ student T-test *p*-value=0.0062. CD8^+^ student T-test *p*-value=0.0002. **(D)** Percentages of PLN CD4^+^ and CD8^+^ T cells from mice in A. **(E)** NOD mice were treated with 3 intravenous peptide injections as before, except that 12-wk pre-diabetic mice were used and scrambled peptide was additionally included as a control. 4 hr after the final injection, PLN WBCs were stimulated with IGRP peptides and brefeldin A for 4 hr, stained for IFNγ and analyzed by flow cytometry as in Figure 3D. Representative animals are shown. **(F)** Mean percentages of cells expressing IFNγ from 6 experiments. IGRP_206-214_ student T-test *p*-value=0.0244. IGRP225-233 student T-test *p*-value=0.0201.

Autoantigen-specific CD8^+^ T cells in the PLNs of prediabetic, OCA-B deficient animals are present but poorly activated by self-peptide (Fig. 3D,E). We treated animals with three injections of JumOCA or scrambled peptide control, and stimulated PLN WBCs with IGRP self-peptides as in Fig. 3D. Significant reductions in IFN expression were observed in JumOCA-treated animals compared to controls (Fig. 7E). Quantification from multiple mice is shown in Fig. 7F. These data show that JumOCA peptide treatment decreases CD8^+^ T cell autoreactivity within the PLNs, similar to genetic OCA-B deletion. Cumulatively, these data provide evidence that targeting OCA-B is a valid strategy to treat emerging T1D, and identify a first-generation inhibitor that is efficacious in vivo.

## Discussion

Autoimmune therapies aim to inhibit autoreactivity while preserving normal immune function. Here we show that genetic and pharmaceutical targeting of OCA-B blocks T1D in mouse models. Although potentially autoreactive CD8^+^ T cell specificities are still generated and appear in the PLNs of OCA-B T cell deficient mice, they fail to become activated and do not accumulate in the pancreas. In contrast, potentially autoreactive CD4^+^ T cells are present but associated with anergic cell populations. OCA-B is not expressed during thymic development or in naïve CD4^+^ T cells, but is stably induced upon activation. In contrast, OCA-B is expressed in naïve CD8^+^ T cells and rapidy increased upon activation.

In CD4^+^ T cells, OCA-B associates with the POU-domain transcription factor Oct1 at ~150 immunomodulatory genes – among them *Il2, Ifng* and *Csf2* (*Gmcsf*). Unlike transcription factors such as NF-AT, AP-1 and NF-kB that act as primary “on” switches, OCA-B removes inhibitory chromatin modifications to establish permissive chromatin environments that allow for silent but previously activated targets to be robustly expressed upon antigen re-encounter (Shakya et al., 2015; Shakya et al., 2011). More specifically, OCA-B interacts with Jmjd1a, a histone lysine demethylase that removes inhibitory histone H3 lysine 9 methyl marks. In vivo, OCA-B loss preserves T cell development and pathogen response, but impairs the establishment of new central memory CD4^+^ T cells. The few cells that are formed respond poorly to antigen re-encounter (Shakya et al., 2015). Repeated antigen encounters that drive high levels of proinflammatory cytokines are a key feature of autoimmunity. In both mice and humans, memory or memory-like cells can underlie autoimmunity including T1D (Chee et al., 2014; Kawakami et al., 2005; Yeo et al., 2018).

The NOD model of T1D is spontaneous but only partially penetrant, much like the human disease (Parker et al., 2009). 60-80% of female NOD mice develop T1D in standard environments (Makino et al., 1985). We find that prediabetic OCA-B T cell deficient NOD mice harbor normal T cell numbers and TCR specificities in their PLNs, consistent with prior observations that OCA-B deficient T cells are largely immunocompetent (Kim et al., 2019; Shakya et al., 2015). Nevertheless, T cell conditional OCA-B knockouts are protected from spontaneous T1D. Whole-body NOD *Ocab* knockouts are also protected from T1D. Prior work showed that OCA-B whole-body knockout protects mice both from lupus-like disease mediated by loss of *Izkf3/Aiolos* (Sun et al., 2003), and in a MOG/EAE model of multiple sclerosis (Ikegami et al., 2019). In contrast, a third study showed exacerbation of systemic, antibody-driven autoimmunity in *Sanroque* mice (Chevrier et al., 2014). The protective vs. exacerbating effects in whole body knockouts may be due to effects on class switching and affinity maturation in germinal centers vs. effects on BCR repertoire (Casellas et al., 2002). A whole body OCA-B knockout also reduces fat accumulation and insulin resistance in aged mice, likely through ablation of B cell-mediated inflammation in white adipose tissue (Carter et al., 2018). T cellspecific knockout of Oct1, a transcription factor with which OCA-B docks, blocks EAE but preserves responses to infection with neurotropic viruses (Kim et al., 2019). Together these data show that in T cells Oct1 and OCA-B promote autoimmunity including T1D, but that their role in B cells may be more complex. OCA-B is typically expressed 50-100-fold higher in B cells compared to T cells. A correctly calibrated dose of competitive inhibitor could therefore be used to selectively blunt T cell-mediated autoimmunity.

scRNAseq experiments reveal decreased pancreatic neutrophil infiltration and an increased percentages of T cells with naïve phenotypes in prediabetic OCA-B conditional knockouts. Gene expression changes were identified consistent with antidiabetogenic effects of OCA-B T cell knockout. For example, the remaining islet neutrophils in knockout mice show reductions in *Cxcl2* expression. Cxcl2 is a potent neutrophil chemoattractant that promotes T1D in vivo (Citro et al., 2015; Diana and Lehuen, 2014). Strikingly, OCA-B T cell deficient mice also lack T1D-associated TCR clones in their activated/memory CD8 pool. Flow cytometry using class I tetramers show a significant reduction in CD8^+^ T cells specific for the IGRP_206-214_ autoantigen. PLN CD8^+^ T cells reactive to this and other autoantigens show reduced reactivity to peptide stimulation ex vivo, suggesting that these autoreactive CD8^+^ cells are produced in the thymus but fail to become activated in the PLNs. In contrast, CD4 clonotypes associated with T1D are present in the pancreata, but associated with anergic gene expression profiles. Consistently, OCA-B loss promotes CD4^+^ T cell anergy in vitro. Loss of Oct1, the transcription factor with which OCA-B docks, also augments anergic responses (Kim et al., 2019).

C57BL/6.RIP-mOVA and NOD.BDC2.5 transgenic mice are simplified monoclonal systems that allow tracking of uniform T cell responses to antigens in vivo (Haskins, 2005). RIP-mOVA expresses synthetic membranous chicken ovalbumin in thymic epithelial and pancreatic beta cells (Kurts et al., 1996; Van Belle et al., 2009). Disease in can be induced in this model in 6-10 d by the addition of OT-I T cells and infection with the OVA-expressing *Listeria*. OCA-B loss in host T cells has no effect in this model. BDC2.5 autoreactive T cells migrate to the islets and cause insulitis beginning at 3-4 weeks (Katz et al., 1993). Due in part to Tregs that have escaped allelic exclusion, in many animals insulitis is limited and does not progress to T1D. However, transferring CD4^+^CD62L^+^Vβ^+^ or CD4^+^CD25^-^ T cells from BDC2.5 donor mice into NOD.SCID recipients also results in rapid (7-10 d) disease onset (Berry and Waldner, 2013). Transplants from both control and OCA-B null NOD/BDC2.5 mice results in disease with similar kinetics. Identical results were generated by depleting CD25^+^ Tregs, or by transplanting total CD4^+^ T cells into immunodeficient NCG recipient mice.

NOD.NY8.3 TCR transgenic mice carry a monoclonal CD8-restricted TCR directed towards IGRP_206-214_ and manifest spontaneous T1D (Verdaguer et al., 1997). Unlike transplantation models, OCA-B T cell conditional NY8.3 mice show significant protection from spontaneous T1D. Cumulatively, the results support a model in which OCA-B loss preserves T cell functionality while desensitizing autoreactive T cells. This results in graded protection that drops off using models in which protection is inseparable from immunodeficiency. These properties make OCA-B a promising target for pharmaceutical inhibition.

Recent work indicates that the prevailing idea that transcription regulators and protein-protein interactions are “undruggable” is erroneous (Antony-Debre et al., 2017; Green, 2016; Skwarczynska and Ottmann, 2015; Zuber et al., 2011). Pre-clinical membrane-penetrating peptides have also been developed that successfully target transcription factors such as BCL6 (Polo et al., 2004). We applied rational-design principles to generate a membrane-permeable competitive peptide inhibitor of the interaction between OCA-B and a downstream effector, Jmjd1a/Kdm3a. The sequence of the inhibitor, termed JumOCA, does not coincide with other sequences in the human or mouse proteome and is not predicted to be strongly immunogenic. JumOCA administration blunts newly-arisen T1D in NOD mice and significantly reduces islet T cell cytokine production. Total T cell numbers were unaffected, however PLN CD8^+^ T cell autoreactivity to IGRP peptides was blunted, similar to T cell knockouts. OCA-B levels in B cells are at least 50-fold higher than in T cells (Heng et al., 2008; Zwilling et al., 1997), making it likely that the observed effects are due to inhibition in T cells. Consistently, three JumOCA injections has no effect of splenic germinal center architecture (not shown), which is known to depend OCA-B expression in B cells (Qin et al., 1998). More than three peptide injections results in toxicity (not shown), complicating experiments testing T1D prevention and durable response. Tat-conjugated peptides are known to be associated with toxicity (Caron et al., 2004; Krautwald et al., 2004; Polo et al., 2004). While these peptides are unlikely to be used in a clinical setting, they offer proof-of-principle for OCA-B as a therapeutic target, and can be used as tools for the further development of therapeutics with the caveat that targeting this pathway would likely inhibit the generation of new memory CD4^+^ T cells.

## Abbreviations

AR: androgen receptor;
DIEA: N,N-diisopropylethylamine;
DCM: dichloromethane;
DMF: dimethylformamide;
HATU: 1-[Bis(dimethylamino)methylene]-1H-1,2,3-triazolo[4,5-b]pyridinium 3-oxid hexafluorophosphate;
ICOS: Inducible T-cell costimulator;
IGRP: islet-specific glucose-6-phosphatase catalytic subunit related protein;
NOD: non-obese diabetic;
PLNs: pancreatic lymph nodes;
PMA: phorbol 12-myristate 13-acetate;
RIP-mOVA: rat insulin promoter-membranous ovalbumin;
scRNAseq: single-cell RNA sequencing;
T1D: Type-1 diabetes;
TFA: trifluoroacetic acid;
TIS: triisopropylsilane;
Treg: regulatory T cell;
UMAP: uniform manifold approximation and projection.

## Author contributions

DT conceived the study and designed experiments, supervised the study, and provided administrative and material support. HK, AS, AI and JLJ acquired and interpreted the data. JP performed analysis of scRNAseq and TCR clonotype data. CG and NPL helped generate critical reagents. BDE, DH-CC, XH and PJ provided material and intellectual support. All authors were involved in writing, reviewing and revising the manuscript.

## Acknowledgements

We thank P. Santamaria, M. Williams, M. Bettini and F. Gounari for critical reading of the manuscript. We thank A. Chervonsky (U. Chicago) for the gift of NOD.CD4-cre mice. We thank J. Marvin and the University of Utah Health Sciences Center Flow Cytometry Core facility for assistance with flow cytometry, and D. Lum and the Preclinical Resource Core for immunodeficient mice. We thank M. Hanson and the University of Utah Health Sciences DNA/peptide synthesis core. This work was supported by grants to DT from the Praespero Foundation, Juvenille Diabetes Research Foundation (1-INO-2018-647-A-N) and National Institutes of Health (R01-AI100873).

## Materials and methods

### Mice

All animal experiments were approved by the University of Utah Institutional Animal Care and Use Committee (IACUC approval 17-05008). NOD/ShiLtJ, NOD.Scid (NOD.Emv302/2.CB17-Prkdcscid), BDC-2.5 TCR transgenic, NY8.3 TCR transgenic and all breeder mice were maintained in a specific pathogen-free research colony. NCG (NOD-Prkdc^em26Cd52^Il2rg^em26Cd22^/NjuCrl) mice were purchased from Charles River Laboratories. NOD.CD4-cre mice were a gift from Alexander Chervonsky (University of Chicago). *Pou2af1*^tm1a(KOMP)Wtsi^ mice were provided by the knockout mouse project (KOMP) repository (University of California, Davis). Mice were derived from embryonic stem cell clone EPD0761_3_B03, generated by the Wellcome Trust Sanger Institute. These cells were derived from C57BL/6N mice, but subsequent mouse crosses were to C57BL/6J (>20 generations) for the RIP-mOVA model and NOD (4 congenic backcrosses plus an additional backcross to NOD.CD4-cre). The embryonic stem cells contain a CSD-targeted allele (Testa et al., 2004). The presence of WT (582 bp) and Post-FLP (700 bp) alleles was determined by PCR using CSD-Pou2af1-F and CSD-Pou2af1-ttR primers. The presence of the null (801 bp) allele was determined using CSD-Pou2af1-F and CSD-Pou2af1-R primers. The presence of the floxed (359 bp) allele was determined using CSD-Lox-F and CSD-Pou2af1-R primers. Primer sequences were as follows: CSD-Pou2af1-F, 5’ TACAGAGAGACTAGACACGGTCTGC; CSD-Pou2af1-R, 5’ GATGAGGACTCTGGGTTCAGAGAGG; CSD-loxF, 5’ GAGATGGCGCAACGCAATTAATG; CSD-Pou2af1-ttR, 5’AGAAGGCCTCGTTACACTCCTATGC.

### Backcrossing *Ocab* germline null and conditional mice to the NOD background

To generate *Ocab^-/-^* and *Ocab* conditional mice on the NOD background, the previously described C57BL/6 *Ocab* germline null allele (Kim et al., 1996) and the newly-generated *Ocab* conditional allele were backcrossed to NOD ShiLt/J mice using congenic markers based on Mouse Genome Informatics (http://www.informatics.jax.org) (Table S1).

### Diabetes development and assessment

Diabetes was monitored using blood glucose test strips (Contour, Bayer). Mice with blood glucose levels >250 mg/dL were considered diabetic.

### Leukocyte isolation and flow cytometry

PLNs were disaggregated and passed through a nylon strainer. Pancreatic leukocytes were isolated as described previously (Sitrin et al., 2013). Briefly, pancreata were chopped and digested using collagenase IV (1 mg/mL, Gibco) in DMEM containing 1% fetal bovine serum (FBS, Rocky Mountain Biologicals) and 10 units of DNase I for 15 min at 37°C. The digested tissues were passed through a 70 μm strainer. Red blood cells were lysed by ACK (Ammonium-Chloride-Potassium) lysis buffer (150 mM NH4Cl, 10 mM KHCO3, 0.1 mM Na2EDTA). For intracellular cytokine staining, cell suspensions in RPMI medium supplemented with 10% FBS were re-stimulated for 4 hr with phorbol 12-myristate 13-acetate (PMA, Sigma, 50ng/mL) and ionomycin (Sigma, 1 μg/mL) in the presence of brefeldin A (GolgiPlug, BD Bioscience, 1 μl/ml), and fixed by cell fixation/permeabilization solution (BD Cytofix/Cytoperm) according to the manufacturer’s protocol. Antibodies used for flow cytometry were as follows: FITC-conjugated anti-mouse CD4, PE-conjugated anti-mouse CD45, PE-conjugated anti-mouse FR4 (BioLegend), PerCP-conjugated anti-mouse CD8a, APC-conjugated anti-mouse IFNγ, PE-conjugated anti-mouse IL-17, PerCP-conjugated anti-mouse CD11b, APC-conjugated anti-mouse F4/80, PE-conjugated anti-mouse Gr-1, PE-conjugated anti-mouse ICOS (eBioscience), PerCP-conjugated anti-mouse FoxP3, V450-conjugated anti-mouse CD73 (BD Bioscience) and FITC-conjugated anti-mouse Vβ8.1/8.2 (Invitrogen). A FACS Canto II (BD Biosciences) was utilized for flow cytometry, and FlowJo software was used for analyses.

### T cell adoptive transfer

6–8 wk old NOD.Scid recipient mice were injected retro-orbitally with 2×10^5^ of splenic CD4^+^CD25^-^ T cells from prediabetic *Ocab^-/-^* or *Ocab*^+/+^ NOD.BDC2.5 donors (6–8 wk old, sex-matched). T cells were purified using a CD4^+^CD25^+^ T cell isolation kit (Miltenyi). For CD4^+^CD25^+^ T cells, 1.5×10^6^ purified splenic CD4^+^ T cells from prediabetic, BDC2.5 transgenic NOD.*Ocab^-/-^* or NOD.*Ocab*^+/+^ mice were transferred to sex-matched 6-8 wk old NCG recipients (University of Utah Preclinical Resource Core) as previously described (Presa et al., 2015). For total splenic transfer experiments, 5×10^6^ splenocytes from prediabetic NOD.*Ocab^fl/fl^* or NOD.*Ocab^fl/fl^*CD4-cre mice were transferred into sex-matched 6-8 wk NOD.Scid recipients mice as described previously (Presa et al., 2015).

### In vitro T cell culture

For anergic cell induction, spleens were harvested from NOD.*Ocab^fl/fl^*CD4-cre or control NOD.*Ocab^fl/fl^* animals. Single-cell suspensions were generated by grinding and passage through 70 μm strainers. Cells were isolated using a mouse CD4^+^ T cell isolation kit (Miltenyi Biotec), and stimulated with 5 μg/ml plate-bound anti-CD3ε (BD Bioscience) ± 2 μg/ml anti-CD28 antibody (eBioscience) as described previously (Kim et al., 2019). For OCA-B inhibitor peptide treatments, cells were isolated from WT C57BL/6 spleens using a mouse naïve CD4^+^ T cell isolation kit (Miltenyi Biotec) and cultured as described previously (Shakya et al., 2011). Indicated concentrations of peptides were incubated with cells, with media changes every 2 days after primary stimulation. Activation of C57BL/6 CD4^+^ cells and subsequent profiling of anergic responses was performed identically to (Kim et al., 2019). For peptide restimulation, total WBCs from PNLs were incubated with 10 μg/mL specific peptide for 4 hr in the presence of brefeldin A. The peptides used were as follows: GAD65_206-214_, TYEIAPVFV; GAD65_546-554_, SYQPLGDKV; IGRP_21-29_, TYYGFLNFM; IGRP_206-214_, VYLKTNVFL; IGRP_225-233_, LRLFGIDLL; IGRP_241-249_, KWCANPDWI; IGRP_324-332_, SFCKSASIP. Cells were then fixed and stained for flow cytometry.

### Single-cell RNA sequencing (GEO Series record GSE145228)

CD3^+^ T cells were isolated from 8 wk NOD.*Ocab^fl/fl^* or NOD.*Ocab^fl/fl^*CD4-cre females using a pan T cell isolation kit (Miltenyi). CD45^+^ pancreatic leukocytes were isolated from 12 wk NOD.*Ocab^fl/fl^* or NOD.*Ocab^fl/fl^*CD4-cre females were isolated by flow cytometry using a FACS Aria (Becton-Dickinson). For each condition, cells were isolated from three mice and combined. Cells were processed using the 10X Genomics Chromium platform according the manufacturer’s instructions. Paired-end high-throughput sequencing (125 cycles) was performed via an Illumina NovaSeq instrument. Sequencing reads were processed by using 10X Genomics CellRanger pipeline and further analyzed using the Seurat R package. Analysis of cells used a standard filtering protocol removing cells with unique feature counts of >4,000 or <500, as well as cells with >5% mitochondrial counts (indicative of dead cells). No more than 15% of total cells were removed by this process. Cells were subjected to unsupervised hierarchical clustering followed by uniform manifold approximation and projection (UMAP) to visualize clusters with similar gene expression and their relative positions.

### Tetramer staining

H-2K^d^ mouse IGRP_206-214_ tetramers (Mouse NRP-V7 mimotope KYNKANVFL) conjugated to APC or PE were synthesized by the NIH tetramer core facility and were a gift of Dr. Maria Bettini (University of Utah).

### OCA-B mutagenesis and Co-IP

For generation of the OCA-B double point mutant, a plasmid transiently expressing human OCA-B, pCATCH-Bob.1 (Gstaiger et al., 1995), was sequentially mutagenized using QuickChange (ThermoFisher). Correct mutagenesis was confirmed by resequencing. HCT116 lysate preparation, co-immunoprecipitation of OCA-B and endogenous Jmjd1a, and immunoblot detection was performed using the same conditions as described (Shakya et al., 2015). For co-IP using M12 cells, lysis buffer consisted of 50 mM Tris-Cl, pH 7.4, 150mM NaCl, 0.5%NP-40, 1 mM EDTA, 1 mM EGTA, plus protease/phosphatase inhibitors (PhosSTOP, Roche). Lysates were incubated with 2.5 μg Jmjd1A antibody (Bethyl) and protein-G Dynabeads (ThermoFisher) in lysis buffer containing 20% glycerol overnight at 4°C. After overnight incubation, the indicated concentrations of peptide or PBS vehicle was added and incubated for a further 3 hr at 4°C prior to bead precipitation and washing 3× in lysis buffer plus 20% glycerol. Co-precipitated OCA-B was analyzed by SDS-PAGE and immunoblot.

### OCA-B inhibitor peptide synthesis

Unique chemically synthesized peptide sequences were as follows: Peptide#1, ARPYQGVRVKEPVK; Peptide#2/JumOCA, VKELLRRKRGH; Peptide#3, ARPYQGVRVKEPVKELLRRKRGH; Scrambled peptide, VLREKGKRHLR. Peptides were synthesized with and without a covalent C-terminal linker and Tat membrane-penetrating peptide (GSG-GRKKRRQRRRGY). Fmoc protected amino acids were obtained from Protein Technologies Inc. 1-[Bis(dimethylamino)methylene]-1H-1,2,3-triazolo[4,5-b]pyridinium 3-oxid hexafluorophosphate (HATU) was purchased from Chemimpex Inc. H-Rink-Amide-ChemMatrix (RAM) was purchased from Biotage. N,N-diisopropylethylamine (DIEA), dichloromethane (DCM), triisopropylsilane (TIS) were purchased from Sigma. Dimethylformamide (DMF), trifluoroacetic acid (TFA), acetonitrile, and ethyl ether were purchased from Fisher Scientific. Peptides were synthesized using automated Fmoc SPPS chemistry (Syro I). Briefly, 220 mg of RAM resin (loading=0.45 mmol/g) was swelled in DCM for 15 min and followed by adding a solution of specific Fmoc protected amino acid (AA = 0.1 mmol, DIEA = 0.2 mmol in DCM = 4 mL) and incubated at room temperature for 1.5 h. The resin was washed with DMF and DCM and incubated with 5 mL of DCM containing 16% v/v MeOH and 8% v/v DIEA for 5 min. This action was repeated 5 times before thoroughly washing with DCM and DMF. Coupling reactions were performed using HATU (5 eq.), DIPEA (10 eq.) and amino acid (5 eq.) in DMF with 15 min heating to 70°C (50°C for Cys). Deprotection was performed using 20% (v/v) piperidine in DMF, 5 min at room temperature for 2 rounds. Peptides were cleaved from the resin by treatment with a cocktail buffer (3 mL/0.1 mmol, TFA:H2O:TIS:EDT = 95:2:2:1) for 2.5 h. Peptide-TFA solution was then filtered and precipitated in cold ether, centrifuged, washed with ether twice and vacuum-dried. The crude product was then purified by RP HPLC. Peptide characterization was performed by LC/MS on an Xbridge C18 5 μm (50 x 2.1 mm) column at 0.4 mL/min with a water/acetonitrile gradient in 0.1% formic acid on an Agilent 6120 Quadrupole LC/MS system. Fractions collected from HPLC runs were also analyzed by LC/MS. The purified fractions containing the targeted product were collected and lyophilized using a Labconco Freeze Dryer. All samples were analyzed by the following conditions: Preparative reverse phase HPLC of crude peptides was performed on a Jupiter 5 μ C18 300 Å (250× 10 mm) column at 3 mL/min with a water/acetonitrile gradient in 0.1% TFA on an Agilent 1260 HPLC system. Purity, isomer co-injection and stability checks were performed on HPLC on Phenomenex Gemini C18 3 μm (110 Å 150×3 mm) column.

### OCA-B inhibitor peptide treatment

Anesthetized prediabetic WT NOD females with glucose levels newly-risen to between 225 to 275 mg/dL were treated with 10 mg/kg inhibitor or control peptides by intravenous (retro-orbital) injection, 3 times every 12 hr. Blood glucose levels were measured after the 2^nd^ and 3^rd^ injection. 4 hr after the last injection, pancreata and PLNs were collected and cell populations were analyzed by flow cytometry.

### Histology

Formalin-fixed pancreatic tissues were embedded in paraffin. H&E-stained sections were scored for islet inflammation based on published precedents (Zhang et al., 2007): 0, sparse surrounding sentinel leukocytes, no insulitis; 1, peri-islet leukocytes; 2, some islet leukocytes, <50% of islet area; 3, islet insulitis with >50% of islet area occupied by leukocytes; 4, islets destroyed with fibrotic remnants.

### Statistical analyses

All error bars denote ±SEM. Two-tailed student T-tests were used to ascribe statistical significance. For all figures, \**p* 0.05; **=*p*≤0.01; ***=*p*≤0.001; ****=*p*≤0.0001.

### Online supplemental material

Fig. S1 outlines the construction and validation of the *Ocab* (*Pou2af1*) conditional mouse allele. Fig. S2 shows results from scRNAseq using PLNs from 8 week-old prediabetic female NOD.*Ocab^fl/fl^*CD4-cre or littermate control NOD.*Ocab^fl/fl^* mice. Fig. S3 shows TCR representation across all populations based on scRNAseq from islet CD45^+^ cells. Fig. S4 shows results from BDC2.5 T cell transfer experiment using OCA-B deficient and sufficient NOD mice, and results from RIP-mOVA T cell transfer experiments using C57BL/6 mice. Fig. S5 shows the concentration of Tat-conjugated peptides in cultured primary CD3^+^ T cells.

## Supplemental information

Figure S1. ***Ocab (Pou2af1)* conditional allele. (A)** Targeting event. Crossing with FLP^Rosa26^ results in conditional (*fl*) allele. Primer-pairs used for genotyping are depicted with arrows. **(B)** Example genotyping of the targeted allele and recombination events. Founder animal is in lane 2. The primer-pairs shown in panel A were used. **(C)** OCA-B immunoblots from spleens of control, *fl/fl* and *Δ/Δ* animals. BJA-B B cell nuclear extract is shown as a positive control (lane 1). **(D)** The *Ocab* (*Pou2af1*) conditional allele was crossed to CD4-cre. Total splenic CD4^+^ T cells were isolated from an *Ocab^fl/fl^*CD4-cre animal or a *Ocab^+/fl^*CD4-cre littermate control, and stimulated for 2 d in vitro using plate-bound CD3ε and soluble CD28 antibodies. An OCA-B immunoblot of stimulated total T cells is shown. β-actin is shown as a loading control. **(E)** To generate NOD.*Ocab* conditional mice by speed congenic backcross, microsatellite repeat polymorphisms at the *Idd* loci were tested by PCR using the primers in Supplementary Table 1. Backcross generations 2 and 4 (B2 and B4) are shown (lanes 3 and 4), together with parent C57BL/6 (lane 1) and target NOD (lane 2) genomic DNA as controls. Images are of different PCR products resolved using agarose or polyacrylamide gel electrophoresis. **(F)** Example genotyping of *Ocab* wild-type and conditional alleles in B4 backcrossed mice. An agarose gel image is shown. Mouse #122 corresponds to the B4 animal shown in B and was used as the founder animal. This animal was crossed NOD.CD4-cre for subsequent experiments with the conditional allele.

Figure S2. **Unaltered T cell populations and TCR representation in prediabetic NOD.*Ocab^fl/fl^*CD4-cre pancreatic lymph nodes compared to NOD.*Ocab^fl/fl^* littermate controls. (A)** CD3^+^ T cells were isolated and combined from the PLNs of three 8 week-old NOD.*Ocab^fl/fl^*CD4-cre mice or three NOD.*Ocab^fl/fl^* littermate controls. scRNAseq was performed. Clusters were overlaid in UMAP plots. **(B)** Six different populations from A were analyzed for differential gene expression. Identified genes are shown as scatter plots. Significantly differentially expressed genes (adjusted *p*-value <0.05) are shown in red. **(C)** % contribution of individual Vb chains is shown. **(D)** *Trbv13-3* expressing cells are shown in blue superimposed on the same UMAP scRNAseq data as in A. The effector CD8^+^ T cell cluster is highlighted in red.

Figure S3. **TCR beta chain representation across the entire population of prediabetic NOD.*Ocab^fl/fl^*CD4-cre pancreatic islets compared to NOD.*Ocab^fl/fl^* littermate controls.** Four 12 week-old mice in each group were combined for scRNAseq TCR analysis.

Figure S4. **Germline and T cell conditional OCA-B loss has no effect in two different T1D transplant model systems. (A)** 5×10^5^ purified CD4^+^CD62L^+^Vb4^+^ T cells from NOD.BDC2.5.*Ocab^-/-^* or heterozygous control NOD.BDC2.5.*Ocab^+/-^* donors were injected retro-orbitally into NOD.SCID recipients (n=7-8 per group). T1D-free survival following transplant is shown. **(B)** 10,000 OT-I T cells were transferred retro-orbitally into C57BL6 RIP-mOVA;*Ocab^fl/fl^*CD4-cre mice or littermate RIP-mOVA;*Ocab^fl/fl^* controls. Mice were infected with 2000 cfu *Listeria monocytogenes* expressing chicken ovalbumin (Lm-OVA) later the same day. Glucose was measured every two days and plotted. Two mice in each group were used.

Figure S5. **Tat-but not OVA-conjugated peptides concentrate in cultured T cells.** CD3^+^ T cells were isolated from the spleens or pancreata of NOD mice and were incubated with 40 μM FITC-conjugated OVA control peptide, or FITC-conjugated to the TAT membrane-penetrating peptide for 15 min. Cells with concentrated FITC fluorescence were observed by epifluorescence microscopy. Images were taken at 40× magnification.

Table S1. **Loci for type 1 diabetes susceptibility and resistance in NOD/ShiLtJ mice, and oligonucleotide primer-pairs used for PCR amplification of *Idd* loci microsatellite markers.** Related to Supplemental Figure S1.

Table S2. **Gene expression comparisons in T cells taken from PLNs of 8 week-old NOD.*Ocab^fl/fl^*CD4-cre and NOD.*Ocab^fl/fl^* littermate controls.** Related to Supplemental Figure S2.

Table S3. **Expression levels of *Ccl1* mRNA in naïve, stimulated, rested and re-stimulated OCA-B/Pou2af1-deficient T cells relative to controls.** Related to Figure 1.

Table S4. **Gene expression comparisons in total CD45^+^ cells taken from pancreatic islets of 12 week-old prediabetic NOD.*Ocab^fl/fl^*CD4-cre and NOD.*Ocab^fl/fl^* littermate controls.** Related to Figure 2.

