## Supplementary figures and images for "Targeting transcriptional coregulator OCA-B/Pou2af1 blocks activated autoreactive T cells in the pancreas and type-1 diabetes"

### Supplemental Figures 1-5

Kim et al. Figure S1

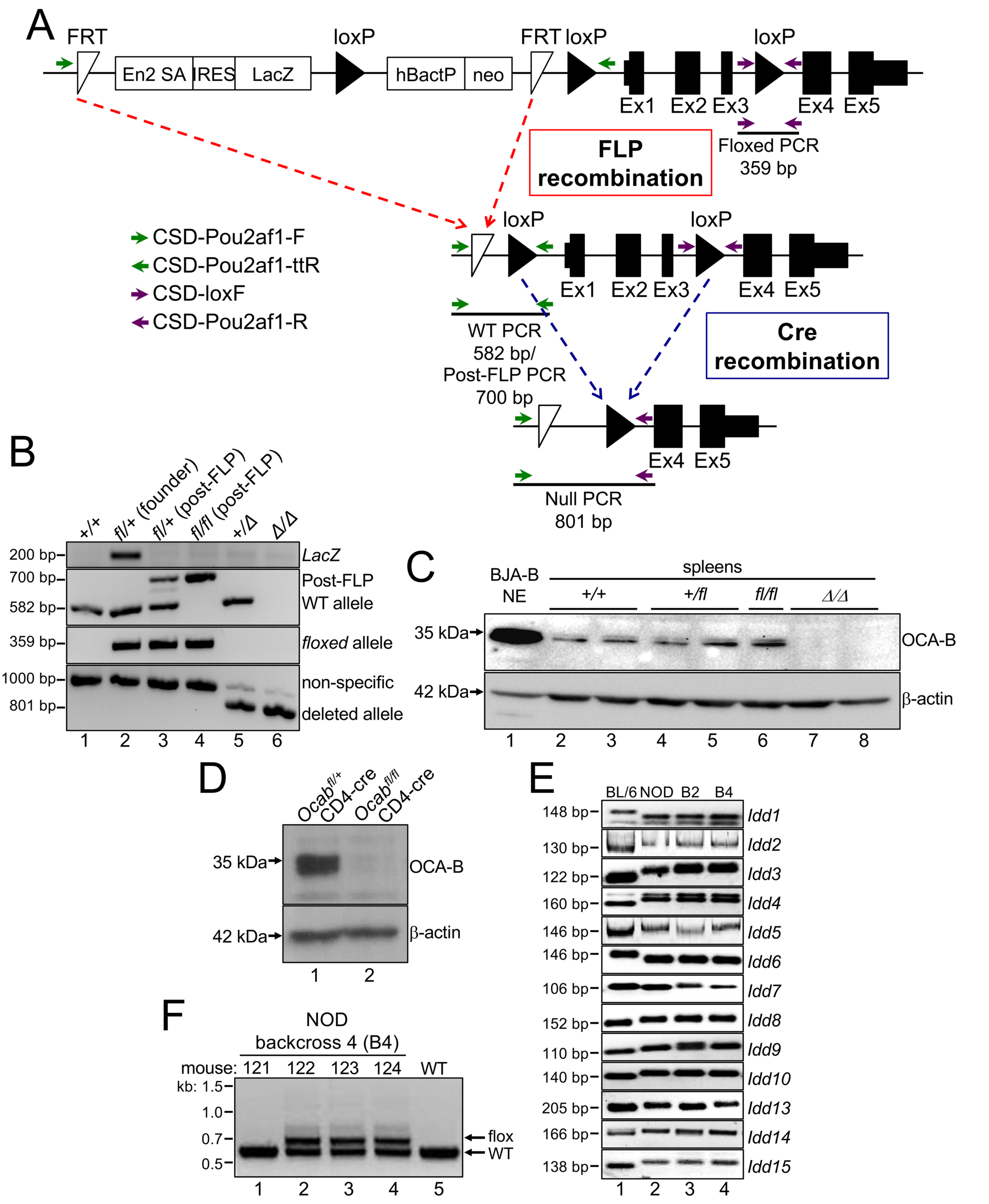

#### Kim et al. Figure S2

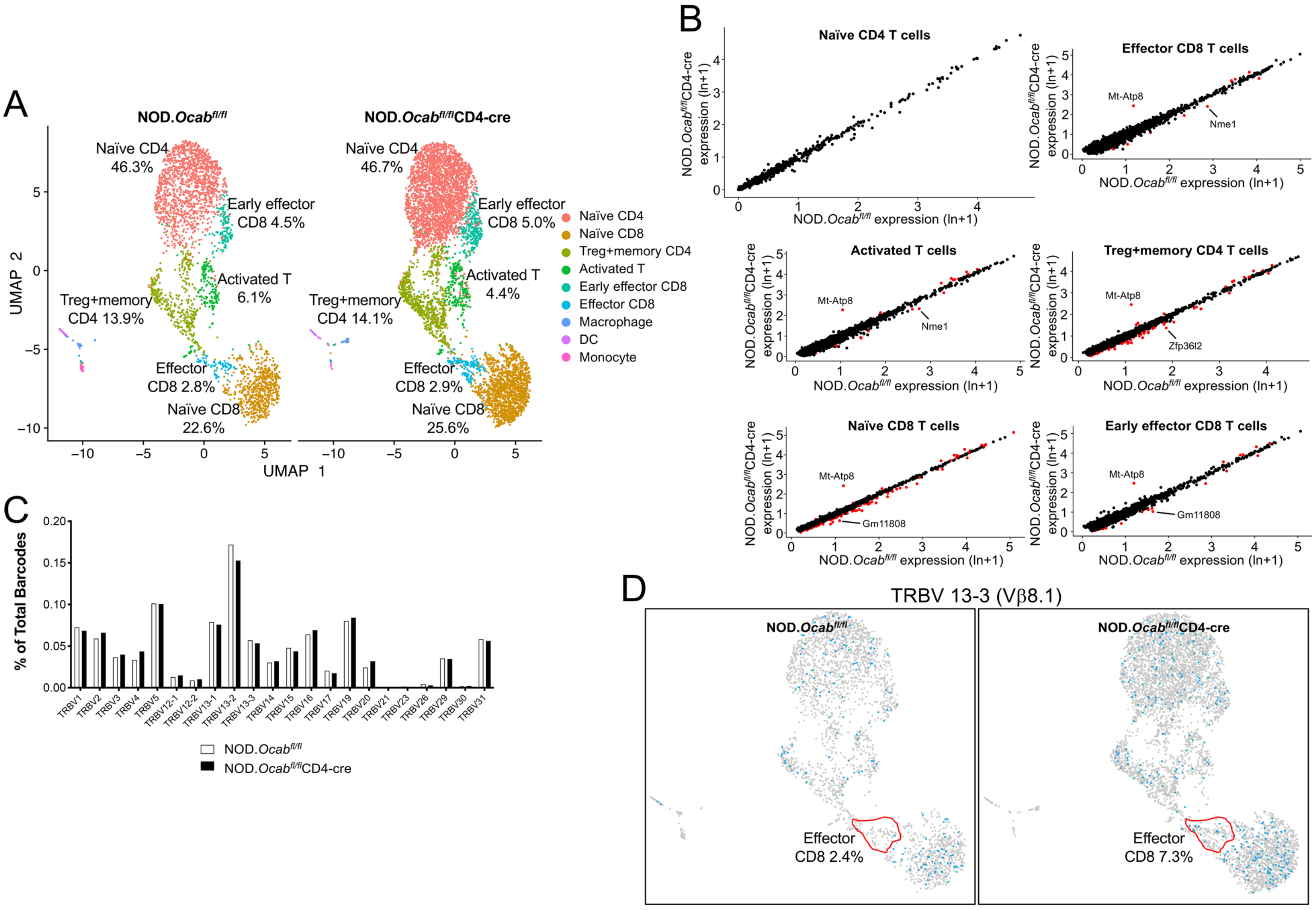

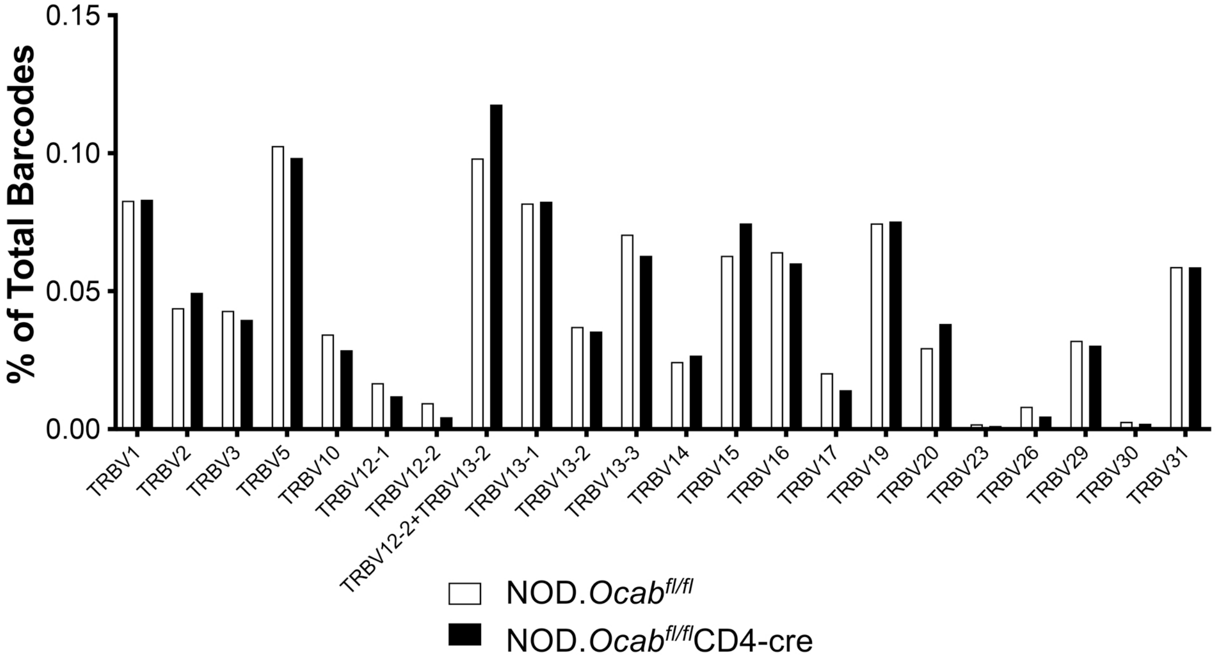

Kim et al. Figure S4

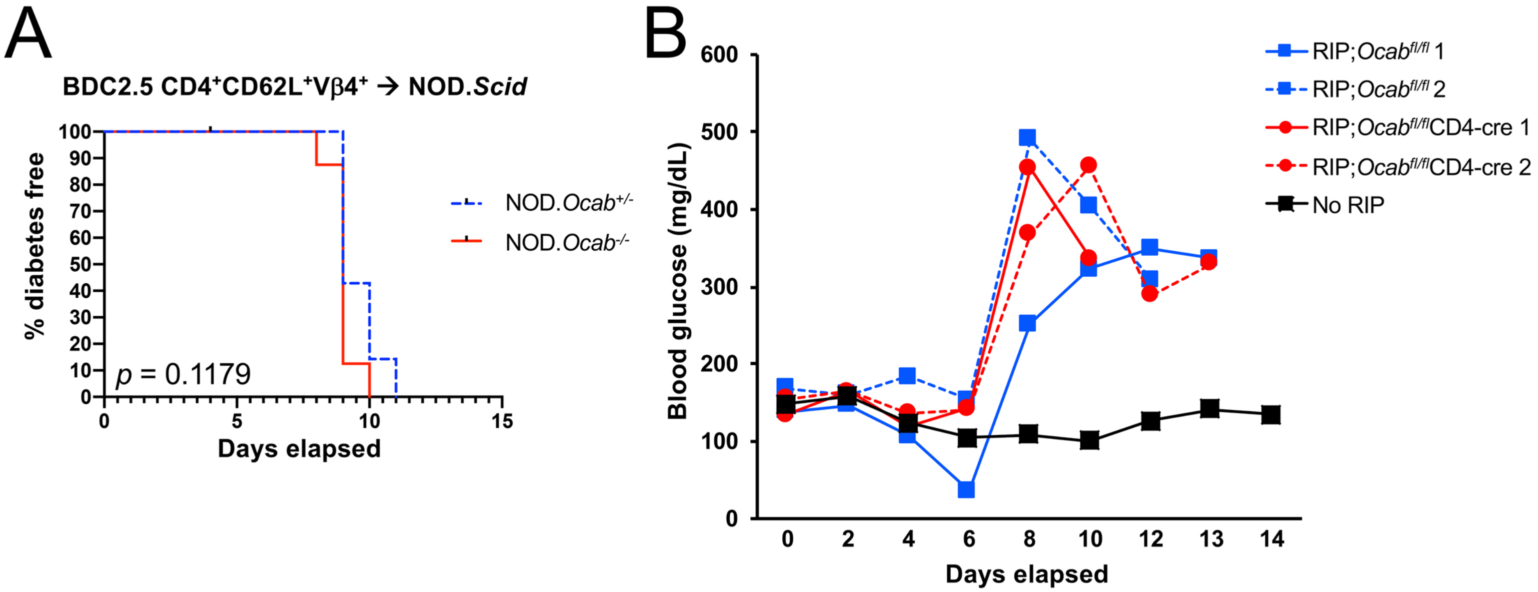

#### Kim et al. Figure S5

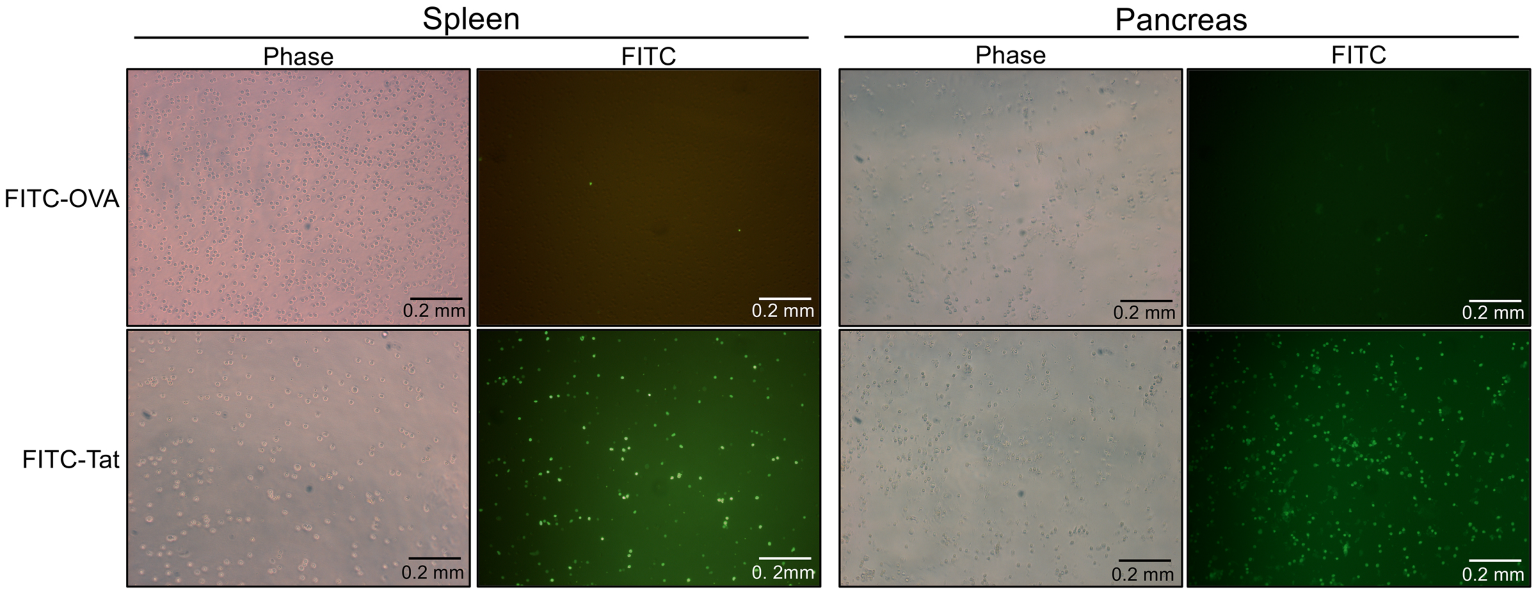
